# A maximum entropy perspective reveals deviations from steady state during active diversification

**DOI:** 10.64898/2026.06.22.733811

**Authors:** Andrew J. Rominger, Katie Thai, Rosemary G. Gillespie, Daniel S. Gruner, John Harte

## Abstract

Ecosystems are rarely at steady state, yet most theory predicting universal biodiversity patterns assumes they are. Here, we test whether and how eco-evolutionary dynamics drive departures from steady state by combining arthropod community data from the geologic chronosequence of the Hawaiian Archipelago with the Maximum Entropy Theory of Ecology (METE), a minimalist steady-state framework that simultaneously predicts species abundance distributions (SADs) and individual metabolic rate distributions (IPDs). The chronosequence of the Hawaiian Archipelago has yielded insights into eco-evolutionary processes because ecosystems growing on different aged substrates offer snapshots of community assembly with different histories. We find that deviations from METE peak at geologically middle-aged sites (150 Kya–1.4 Mya), consistent with active adaptive radiation pushing communities away from statistical steady state. Within-site *β*-diversity, which also peaks at middle-aged sites, robustly predicts deviations from METE across all sites, while the proportion of non-native species predicts deviations only after excluding the geologically youngest site. Partitioning *β*-diversity between native and non-native species resolves this discrepancy: at the youngest site, non-native species are distributed homogeneously and do not elevate *β*-diversity despite their high proportional representation. Together, these results are consistent with a trajectory from young, dispersal-assembled communities near statistical steady state, through an eco-evolutionary non-steady-state transition driven by diversification, to a new stable steady state at the oldest sites. Our findings suggest that periods of active diversification create windows of ecological instability that may facilitate biological invasion, with implications for understanding invasion dynamics in biodiversity hotspots.

## Introduction

Biodiversity patterns are bound by two seemingly contradictory states: they show striking consistency in form across systems, and yet also must emerge in spite of, or because of, the constant flux and dynamic change of ecology and evolution. The near universality of biodiversity patterns includes the scaling of species richness with area (Harte *et al*. 2009; Rosenzweig *et al*. 1996), the ubiquity of rare species and scarcity of common species (Harte 2011; Hubbell 2001), and the highly skewed distribution of biomass or metabolic resource use across organisms (Brown 1995). Debate has persisted for over a century as to whether these consistent patterns form because of ecological and evolutionary drivers that force systems into these configurations, or are largely the outcome of chance acting on large systems of constituent parts (Adler *et al*. 2007; Arrhenius 1921; Chisholm & Pacala 2010; Clements 1916; Diaz *et al*. 2021; Fisher *et al*. 1943; Gleason 1926; Harte 2011; Hubbell 2001; Hughes 1986; MacArthur & Wilson 1967; Magurran 2005; Preston 1948; Rosenzweig *et al*. 1996). The deterministic view gives rise to niche-based theories of coexistence (Barabás *et al*. 2018; Chesson 2000), while the stochastic view has produced a number of neutral-like models (Harte 2011; Hubbell 2001).

Both deterministic and stochastic perspectives on the generation of universal biodiversity patterns most often assume an ecosystem is at “equilibrium” or steady state (Barabás *et al*. 2018; Chesson 2000; Harte 2011; Hubbell 2001). However, ecosystems are known to be dynamic both through intrinsic ecoevolutionary processes (Gillespie 2004; Knope *et al*. 2012; Rominger *et al*. 2016) and due to recent anthropogenic perturbation (Barton *et al*. 2021; Chen *et al*. 2011; Dobrowski *et al*. 2013; Garcia *et al*. 2022; Graham *et al*. 2017; Lenoir & Svenning 2015). Whether universal patterns are insensitive to non-steady-state processes or are a result of those processes remains un-addressed. To better understand the interaction of non-steady-state processes and universal patterns, we combine data (Gruner 2007) from the highly dynamic forest ecosystems of the Hawaiian Archipelago (Wagner & Funk 1995) with the intentionally minimalist steady state predictions of the Maximum Entropy Theory of Ecology (Harte 2011).

### The Hawaiian Archipelago Chronosequence

The Hawaiian Archipelago holds numerous examples of adaptive radiation (Baldwin & Sander-son 1998; Bennett & O’Grady 2013; Gillespie 2004; Givnish *et al*. 2009; Knope *et al*. 2012; Lerner *et al*. 2011; Magnacca & Price 2015). Those remarkable examples of diversification along with the geographic isolation and chronological geology of the archipelago have made it a focal point for the study of eco-evolutionary processes (Barton *et al*. 2021; Chadwick *et al*. 1999; Gillespie & Clague 2009; Price & Wagner 2004; Simon 1987; Wagner & Funk 1995). The manner in which the islands formed in chronological sequence Figure 1 has allowed for the study of ecosystems in different stages of eco-evolutionary assembly (Gillespie 2004; Rominger *et al*. 2016; Vitousek 2004) by considering ecosystems on different aged substrates to be snapshots of eco-evolutionary processes at different time points.

**Figure 1:**
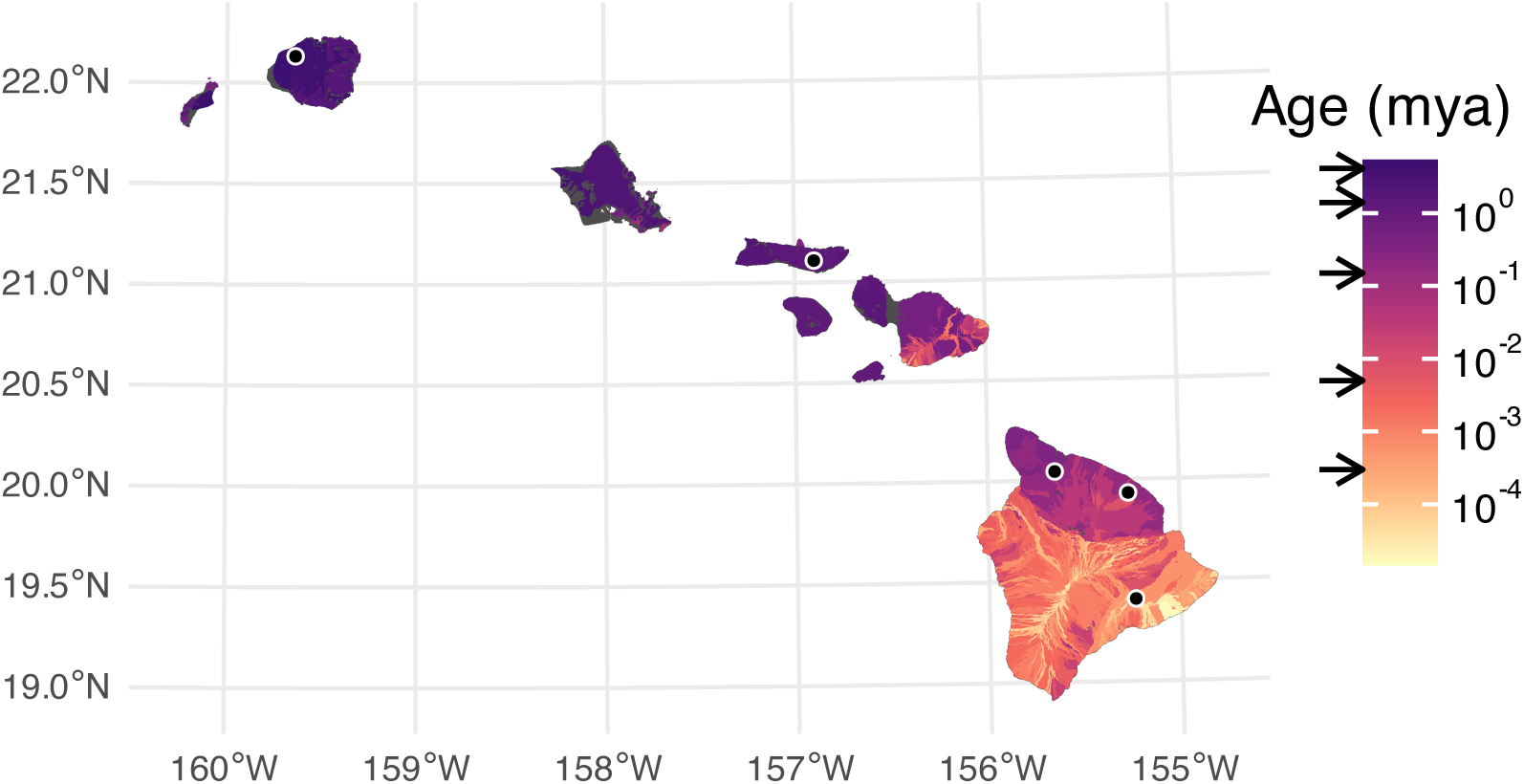
Chronosequence of the Hawaiian Archipelago showing the oldest flows to the north west on Kaua‘i and youngest flows to the south east on Hawai‘i. Substrate age data from Sherrod *et al*. (2021). The five sites are indicated with dots on the map and arrows indicating their flow ages on the legend. From oldest to youngest sites are: Alaka‘i (Kaua‘i: 4.1 Mya), Kamakou (Moloka‘i: 1.4 Mya), Kohala, Laupāhoehoe, Kīlauea (Hawai‘i: 0.5 Mya, 150 Kya, 300 ya).

The Hawaiian Archipelago has also absorbed pervasive anthropogenic changes to its biodiversity, with a rapidly expanded human land-use footprint and markedly accelerated introduction of non-native species following Western contact and settler colonialism (Gon *et al*. 2018; Graham *et al*. 2017). As such the Hawaiian Archipelago has produced numerous studies of rare and endangered species (Magnacca 2007; Pratt *et al*. 2009; Régnier *et al*. 2015; Sakai *et al*. 2002) and the functions of non-native species in ecosystems (Barton *et al*. 2021; Ostertag *et al*. 2009).

As a result of the combination of ancient and recent processes, arthropods assemblages across the Hawaiian Archipelago are now composed of members of adaptive radiations (Bennett & O’Grady 2013; Gillespie 2004; Magnacca & Price 2015), many rare species (Magnacca 2007; Magnacca & Price 2015), and introduced species (Nishida 1992). Given the abundance and diversity of arthropods, they also present a unique opportunity to use ecological theory to study how biodiversity conforms to or deviates from steady state.

### The Maximum Entropy Theory of Ecology

The Maximum Entropy Theory of Ecology (METE Harte (2011)) is a particularly useful tool for studying steady state biodiversity patterns because it produces multiple testable predictions, including the distribution of abundances across species and the distribution of metabolic rates across individuals, making falsifications of the theory more informative (McGill 2003). The species abundance distribution (SAD) and distribution of metabolic rates across individuals (IPD for individual power distribution; power because metabolism is energy per time) contain complementary and overlapping information about an assemblage. The SAD contains information about species diversity and encodes signals of different demographic processes (Engen & Lande 1996b, a). The IPD contains information about how energetic resources are partitioned across individuals which encodes information both about the life history strategies of different lineages and the efficiency with which organisms can acquire useful resources from their environments for reproduction (Brown *et al*. 1993; Burger *et al*. 2019). This link to reproduction interconnects abundance and metabolic rate distributions (Brown *et al*. 2004; Damuth 1987).

METE arrives at some similar predictions to other null or neutral models of biodiversity such at the Unified Neutral Theory (Hubbell 2001) but instead of being limited to a single neutral formulation, METE is grounded in the principles of statistical mechanics Jaynes (1957) drawing from the probabilistic properties of large, randomly assembled systems. METE posits that biodiversity patterns are statistical outcomes from a few simple state variables, independently of the specific biological mechanisms governing the values of the state variables. The summed metabolic rate across all individuals (Harte 2011). Thus, its predictions represent a community in statistical steady state. The steady state assumption holds so long as the governing biological mechanisms do not introduce information beyond what the state variables themselves capture. Examples of such additional information could be rapid environmental change leading to mismatched niches with local environments, rapid diversification, or biological invasion. Under such scenarios the simple state variables of *S*_0_, *N*_0_, and *E*_0_ would not likely constitute enough information to yield accurate predictions for broad biodiversity patterns such as the species abundance distribution or distribution of metabolic rates across individuals. Thus finding deviations between observed patterns of abundance and metabolic rate distributions compared to their METE predictions allows us to identify ecological systems out of statistical steady state (Harte 2011; Newman *et al*. 2020; Rominger *et al*. 2016).

### METE across the chronosequence

In this study of arthropods communities across the chronosequence, we find consistent deviations from steady state at middle-aged sites. We therefore ask: what could be the drivers of non-steady state at these middle-aged sites? Our motivating hypothesis is that steady state is reached quickly at the youngest site through dispersal-based assembly, transitioning to non-steady state assembly at middle-aged sites driven by evolutionary diversification that culminates at the oldest site in a new steady state. With the data at hand we can begin to chip away at that larger hypothesis by investigating two corollary hypotheses: 1) that peak diversification at middle aged sites leaves a signature of increased *β*-diversity (Economo & from METE; and 2) the spread of non-native species, an inevitable confounding factor in the Hawaiian Archipelago, disrupts eco-evolutionary assembly and results in non-steady-state and thus departures from METE.

Deviations from METE are indeed predicted by within site *β*-diversity which peaks at middle aged sites, consistent with rapid diversification processes. Deviations are also partially predicted by proportion of non-native species at a site, except that the proportion of non-native species is high at the youngest site while this site also shows some of the lowest deviations from METE. The conundrum of the youngest site is resolved by considering the contributions of native and non-native species to *β*-diversity. The high contribution of native species to *β*-diversity at middle-aged sites again supports the possibility of rapid diversification playing a role in departures from steady state. More work connecting deviations from steady state to ecological and evolutionary drivers will be needed to further test these possibilities.

## Methods

### Reproducibility and data preparation

A custom R package, *HawaiiArthMETE* (Rominger & Thai 2026), accompanies this paper and can be used to reproduce all results. All data analyzed here are available via the package as described in Supplementary Section S1.

Arthropod data come from the original publication by Gruner (2007), a study that collected 17024 specimens across the chronosequence Figure 1. The five study sites in Gruner (2007) were standardized to be climatically homogeneous (Vitousek 2004), seeking to capture differences in arthropod assemblages due to substrate age and not other abiotic factors. Arthropod assembledges were collected by canopy fogging from the endemic ‘ōhi‘a lehua (*Metrosideros polymorpha* Gaud. Myrtaceae), which is the dominant tree in mesic to wet, mid-elevation forests the Hawaiian Archipelago. Each site contained between 8 and 11 replicate samples.

Each specimen was identified to the species level (morpho species were used in 31.56% of specimens). Specimen lengths were measured to the nearest millimeter. For METE analyses we need to estimate metabolic rate from these data. We use a 3/4 scaling rule (Brown *et al*. 2004) to calculate metabolic rate as *B* = *M*^3/4^ where is mass. Because METE normalizes metabolic rates such that the lowest possible rate is 1 we can safely ignore units and any scaling constants (e.g. *b*_0_ in *B* = *b*_0_*M*^3/4^).

This leaves the estimation of mass from length. Gruner (2007) used taxon– and stage-specific length-to-biomass allometric scaling relationships (Gruner 2003) to approximate mass as the expected value of a log-log regression

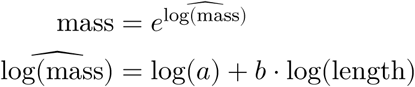

where *a* and *b* are estimated from data collected by Gruner (2003) and vary by taxon and stage.

Using these expected value masses based on length, the latter measured to the nearest millimeter, yields metabolic rate approximations that are artificially discretized (see the appendix “Sampling biomass estimates from allometric equations”). These discretizations in turn lead to spurious conclusions about the predictiveness of METE because METE assumes metabolic rates are continuous (see Supplementary Section S6).

Therefore, instead of estimating mass as the expected value from an allometric relationship we sample masses as random draws from the full model relating mass to length

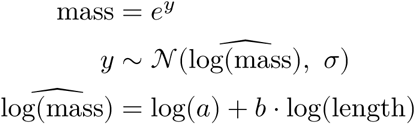

where σ is the taxon– and stage-specific standard deviation of residuals. The full procedure for sampling masses is detailed in the appendix “Sampling biomass estimates from allometric equations” and the consequences of using random draws versus expected values on the METE analysis is thoroughly investigated in Supplementary Section S6. We conclude that using random draws for mass estimates is largely similar to using expected values, and our overall conclusions are unchanged, but using draws avoids both spurious deviations from and conformations to METE that would be introduced by using expected values of mass.

### METE analysis

We produce METE predictions for each sample within a site, including both native and non-native species in the calcualtion of state variables, SADs, and IPDs. We use the *meteR* R package (Rominger & Merow 2017) for all METE calculations. We compare METE predictions to observed SAD and IPD patterns using a *z*^2^-value. The use of *z*^2^-values is discussed in Rominger & Merow (2017), but briefly: the *z*^2^-value compares the log likelihood of the data given the METE model to the log likelihood of a hypothetical data set drawn from the METE model itself. This comparison is done many times and the actual *z*^2^-value is calculated as

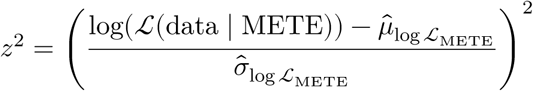

where 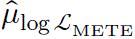 is the mean of the log likelihood of data drawn from METE itself, and 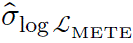 is the standard deviation of the log likelihood of data drawn from METE itself.

If METE is truly correct in describing the data, *z*^2^ will asymptotically have a *χ*^2^ distribution with *d*.*f*. = 1 (Rominger & Merow 2017). If METE does not truly describe the data, the *z*^2^-value is likely to be large.

We also quantify the difference between METE predictions and observed SADs and IPDs by comparing the tails of the observed and predicted distributions. These tails can be especially informative about the nature of deviations (Harte 2011). For the SAD we compare rarity as the log ratio of observed proportion of singletons to the predicted proportion of singletons, and we compare dominance as log ratio of observed total abundance of the top three most abundant species to the predicted total abundance of the top three most abundant species. For the IPD we compare metrics similar to rarity and dominance: the log ratio of observed proportion of individuals with a scaled metabolism of less than 3 to its prediction, and the log ratio of the observed sum of metabolic rates for the top 10 individuals to its prediction.

### Statistical analysis of deviations from METE and potential explanatory variables

We first evaluate how deviations from METE, as measured by *z*^2^-values, vary across the chronosequence. We then test for relationships between *z*^2^-values and proportion of non-native species at a site as well as within site β-diversity.

For all these analyses we fit linear models predicting log-transformed *z*^2^-values responding to each explanatory variable: substrate age, proportion non-native species, and within site β-diversity. Log-transforming *z*^2^-values brings them closer to the normal assumptions of a linear model. The need for log-transformation is not surprising: while *z*^2^-values will be *χ*^2^distributed if METE is true, deviations from METE add more probability to larger values, and taken together will lead to a right-skewed distribution of *z*^2^-values. Log-transformation brings right-skewed random variables closer to normal.

When modeling deviations from METE, we have two statistical challenges to address: samples within sites could represent pseudo replicates necessitating a random-effects model; however, because explanatory variables are site-level characteristics, site-level random effects would be completely collinear with site-level explanatory variables.

To address these challenges we model the site-level mean and variance rather than each individual sample within each site. This is the most conservative approach but runs the risk of being under-powered because while we have 48 samples, we only have 5 sites. Modeling mean and variance is the same technique used in meta-analyses. We therefore make use of the *metafor* package (Viechtbauer 2010) in fitting and testing these models.

### Calculating ***β***-diversity

To quantify within site *β*-diversity as the mean dissimilarity between samples within a site. We first calculate mean Bray-Curtis dissimilarity between samples and then, to standardize *β*-diversity measures across sites, we normalize mean dissimilarity to a null model. We use the quasiswap_count algorithm implemented in *vegan* (Oksanen *et al*. 2025) to generate null samples. This algorithm preserves row and column sums of a sample by species matrix and also seeks to maintain null matrix fill close to the observed fill thereby mimicking patchiness in the real data. Within each site we use 500 null sample-by-species matrices to generate null models for each site. The final measure of within site *β*-diversity is then calculated as a z-score

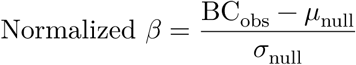

where *BC*_obs_ is the observed mean Bray-Curtis dissimilarity between samples within a site, *μ*_null_ is the mean of the null distribution, and *σ*_null_ is the standard deviation of the null distribution.

We are also interested in understanding how native and non-native species contribute to the overall mean dissimilarity at a site. We can do this by partitioning Bray-Curtis dissimilarity between native and non-native groups of species. The end result of partitioning Bray-Curtis is two dissimilarity values (one for native species, one for non-natives) that add to the total dissimilarity at a site. Supplementary Section S5.3 provides the mathematical and computational details of how we do this.

We also compare this Bray-Curtis dissimilarity partitioning to a null model. We again use the quasiswap_count algorithm but must constrain the algorithm to permute only inside submatrices (one for native species, another for non-natives). This additional constraint is to ensure that the distributions of native and non-native abundances across sites are preserved in permuted and observed site-by-species matrices. We wrote custom code to do this in the *HawaiiArthMETE* package and described in Supplementary Section S5.3.

Comparing observed Bray-Curtis dissimilarity partitions to their permutational null counterparts yields a z-score for the partitioned dissimilarities. While partitions of Bray-Curtis dissimilarity perfectly add to equal the overall dissimilarity, these z-scores will not perfectly add to the previously described overall dissimilarity z-scores because the partitioned z-scores are calculated from a null model with additional constraints (considering native and non-native species) compared to the overall z-scores.

## Results

### Deviations from METE across the chronosequence

Deviations from METE in the SAD and IPD are largely in agreement: samples with high *z*^2^-values for the SAD tend to have high *z*^2^-values for the IPD (see Supplementary Figure S2). The *z*^2^-values for both distributions peak at middle aged sites along the chronosequence Figure 2. For the SAD this is supported by a likelihood ratio test comparing a quadratic model of *z*^2^-values to an intercept-only model with LRT statistic Λ = 28.73, *d*.*f*. = 2, *P* < 0.0001.

**Figure 2:**
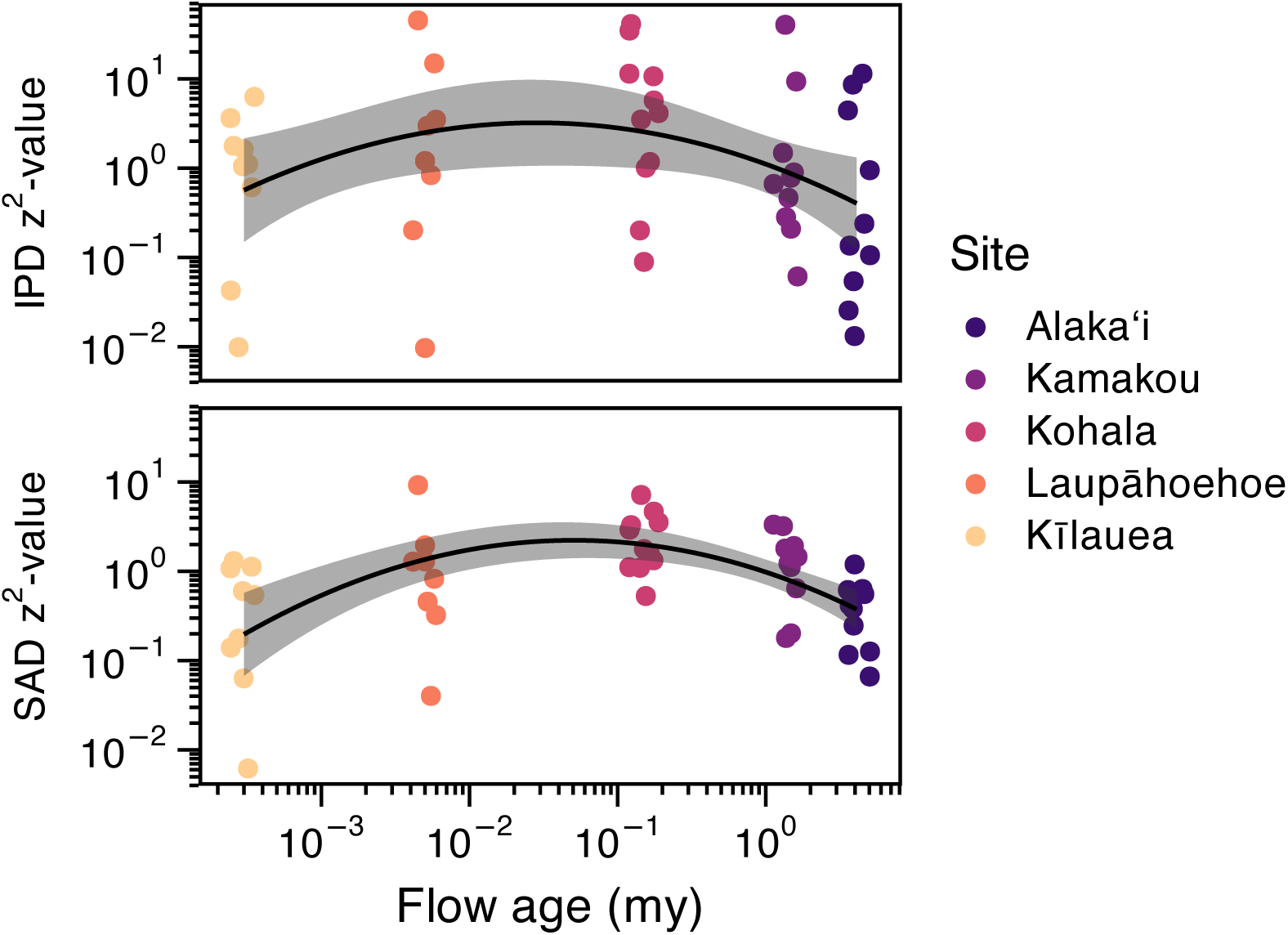
Deviations from METE across the chronosequence as measured by *z*^2^-values. Colors represent sites and match the color coding of substrate age in Figure 1.

The overall spread of IPD *z*^2^-values is greater than that of SAD *z*^2^-values. Not surprisingly, for quadratic model of IPD *z*^2^-values across the chronosequence, the LRT statistic is Λ = 5.513, *d*.*f*. = 2, *P* = 0.0635. While the *P*-value for the test of the quadratic model for IPD *z*^2^-values just misses the ≤ 0.05 significance cut off by the likelihood ratio test, the less conservative Wald test does show the quadratic term to be significant with a 95% confidence interval of [–0.154, –0.014], which does not overlap 0.

Figure 3 shows two exemplar SADs and IPDs with low and high deviation from METE predictions. For the SAD, the primary driver of differences between METE predictions and observed patterns is that observed species abundances have both a higher proportion of singletons and a larger concentration of abundance in the most abundant species Figure 4 compared to METE. Taken together, this means that observed SADs are more uneven than those predicted by METE.

**Figure 3:**
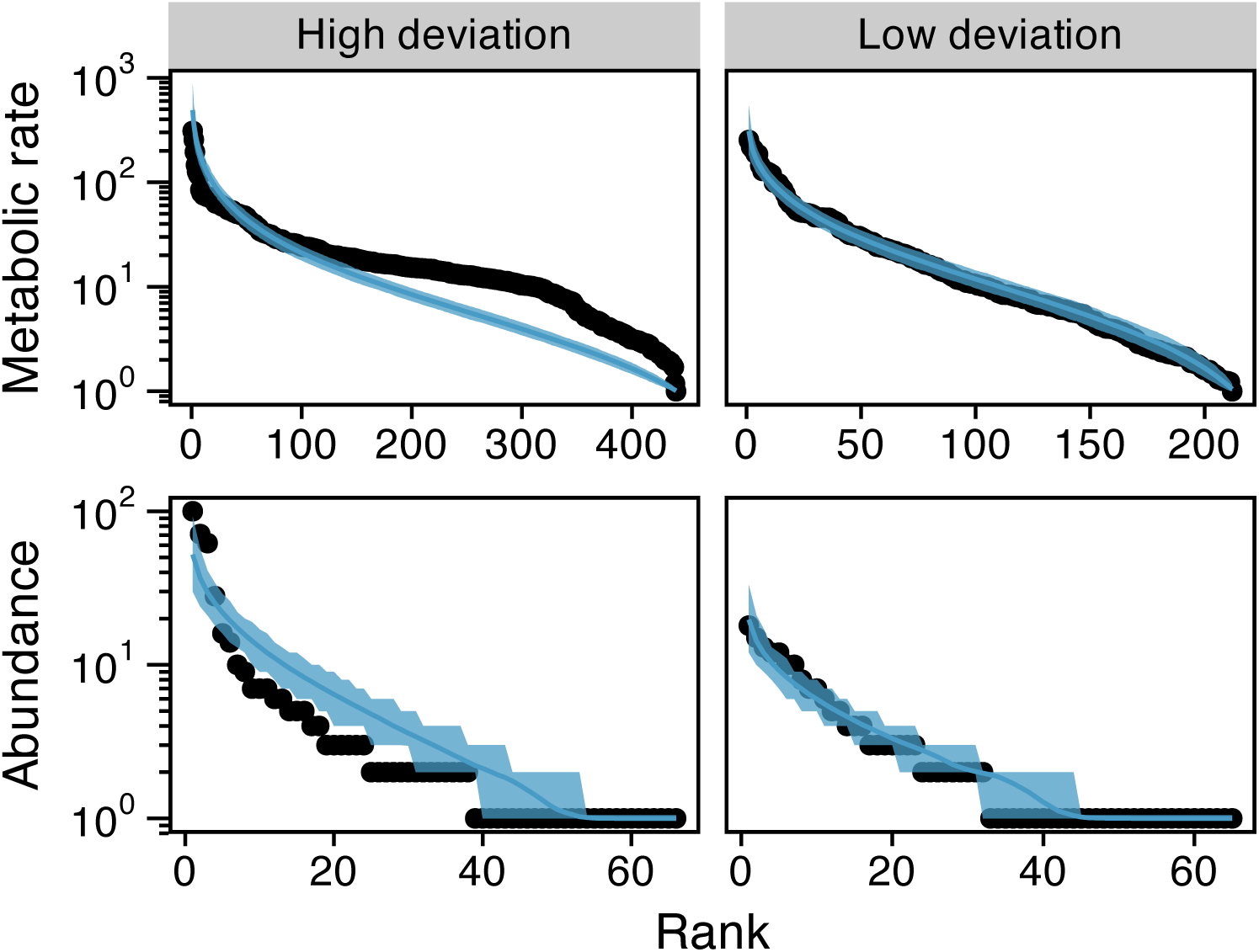
Two exemplar IPDs and SADs showing high and low deviation from METE. The low deviation distributions come from the same sample at the Alaka‘i site while the high deviation distributions come from the same sample at the Kamakou site. The black dots show the observed data while the blue line and shadded blue region show the central tendency and 95% confidence envelopes of 1,000 random draws of data from the SAD and IPD predicted by METE.

**Figure 4:**
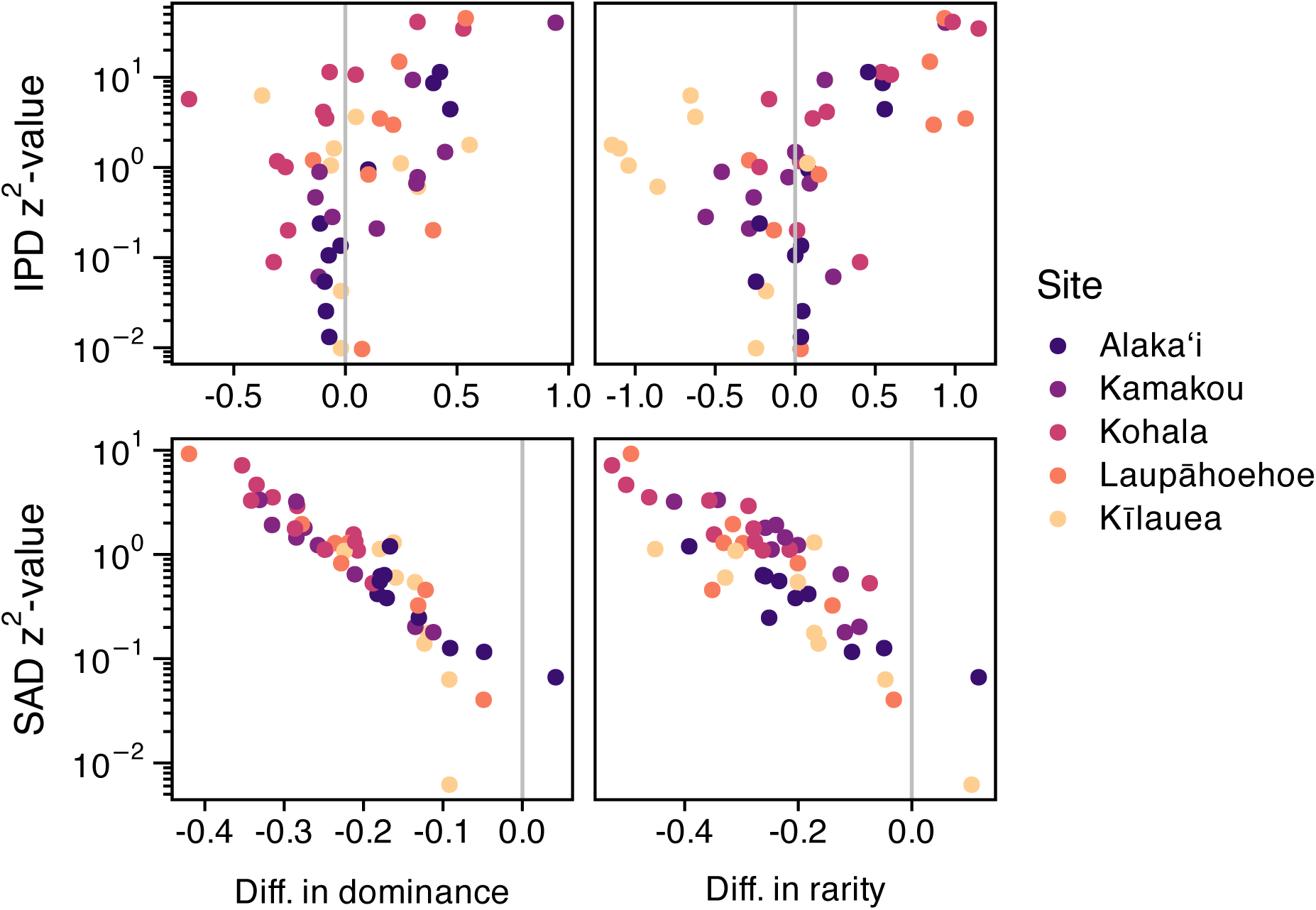
Relationship between *z*^2^ measure of deviation from METE and different metrics describing how observed data and METE predictions diverge.

For the IPD there is not a consistent relationship between deviations and the tails of the metabolic rate distribution Figure 4. In terms of proportion of individuals with very low metabolisms, 19 samples showed theory over-predicting this proportion while 27 samples showed theory under-predicting this proportion. The difference was not significant by a binomial generalized linear model of over-versus-under prediction grouped by site (z = –1.173, = 0.241). In terms of concentration of metabolic rate in the most metabolically active individuals, 24 samples showed theory over-predicting this concentration while 24 samples showed theory under-predicting this concentration. With the two numbers exactly equal, any *P*-value will never be significant.

### Deviations from METE predicted by non-native species and ***β***-diversity

The proportion of non-native species at a site has a significant linear relationship with both SAD and IPD *z*^2^-values when excluding Kīlauea Figure 5. For the SAD the LRT of the linear model with positive slope versus the intercept only model is Λ = 22.329, *d*.*f*. = 1, *P* < 0.0001. For the IPD the LRT of the linear model with positive slope versus the intercept only model is Λ = 5.262, *d*.*f*. = 1, *P* = 0.0218.

**Figure 5:**
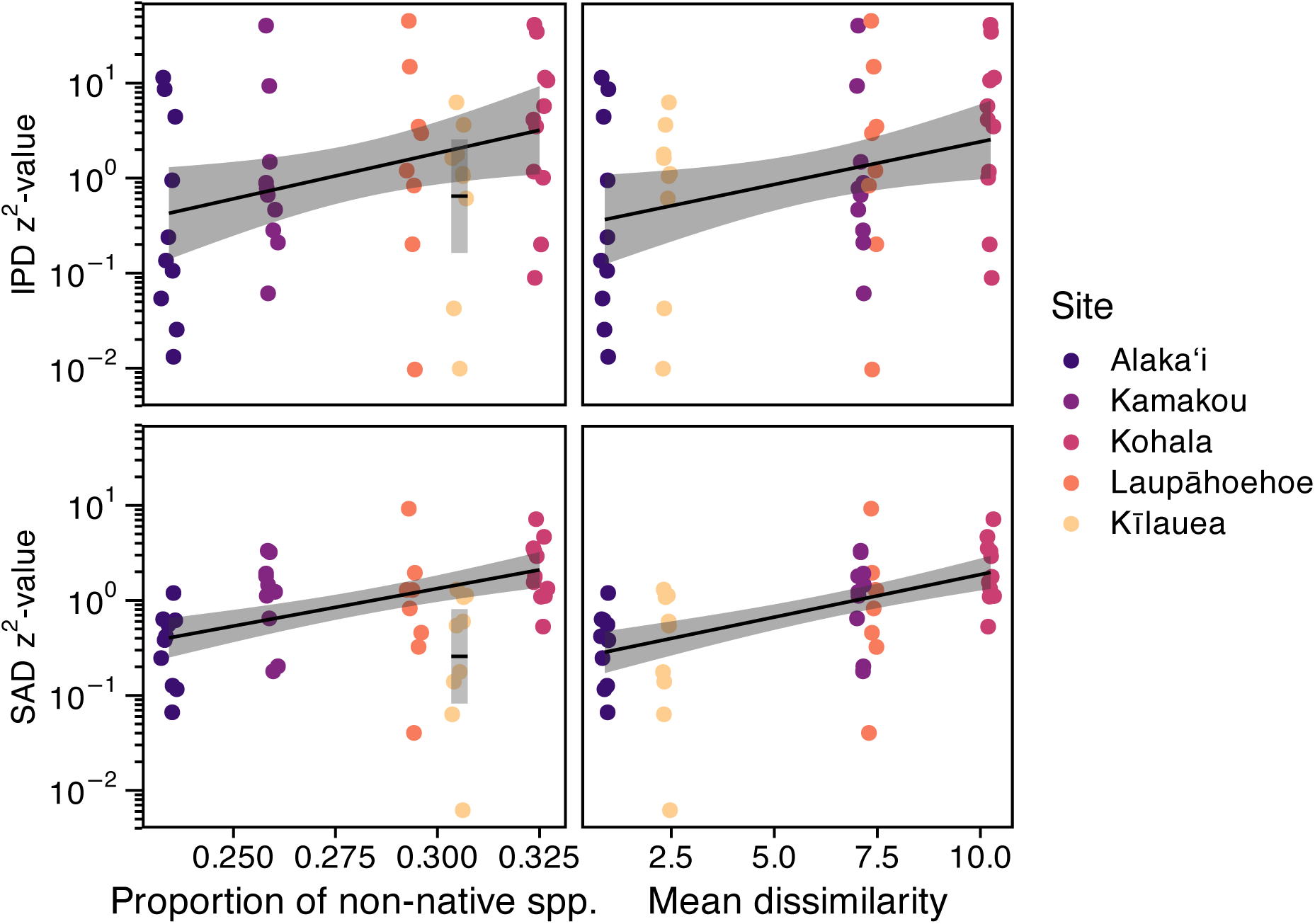
Relationship between *z*^2^ measure of deviation from METE and different predictor variables: proportion of non-native species and β-diversity.

Kīlauea is an interesting outlier in both the SAD and IPD *z*^2^-values relationships with proportion non-native species. In the case of the SAD, the 95% confidence interval for the mean of Kīlauea *z*^2^-values is well below the 95% confidence envelope of the overall relationship Figure 5. In the case of the IPD, the 95% confidence interval of the mean of Kīlauea *z*^2^-values is below the 95% confidence envelope of the overall relationship, but the confidence interval and envelope do partially overlap Figure 5. This is a strong suggestion that the effect of non-native species at Kīlauea is distinct from other sites. We unpack that possibility further in our analysis of *β*-diversity.

Within site *β*-diversity has a significant positive linear relationship with the *z*^2^-values of both the SAD and IPD. For the SAD the LRT of the linear model with positive slope versus the intercept only model is Λ = 29.968, *d*.*f*. = 1, *P* < 0.0001. For the IPD the LRT of the linear model with positive slope versus the intercept only model is Λ = 5.471, *d*.*f*. = 1, *P* = 0.0193. There are no clear outlier sites in the relationship of *z*^2^-values and β-diversity.

### Partitioning *β*-diversity across native and non-native species

Kīlauea is an outlier in the relationship between proportion of non-native species and *z*^2^-values but not an outlier in the relationship between β-diversity and *z*^2^-values. To help understand what is different about Kīlauea we partition native and non-native species’ contributions to *β*-diversity. We find that while Kīlauea has a high proportion of non-native species, it also has a low β-diversity, and non-native species contribute negatively, compared to the null, to that low *β*-diversity Figure 6. Taking these insights together, Kīlauea has many non-native species, but those species are distributed very homogeneously across samples. The oldest site Alaka‘i reverses this pattern: native species contribute negatively to the overall (low) *β*-diversity at that site.

**Figure 6:**
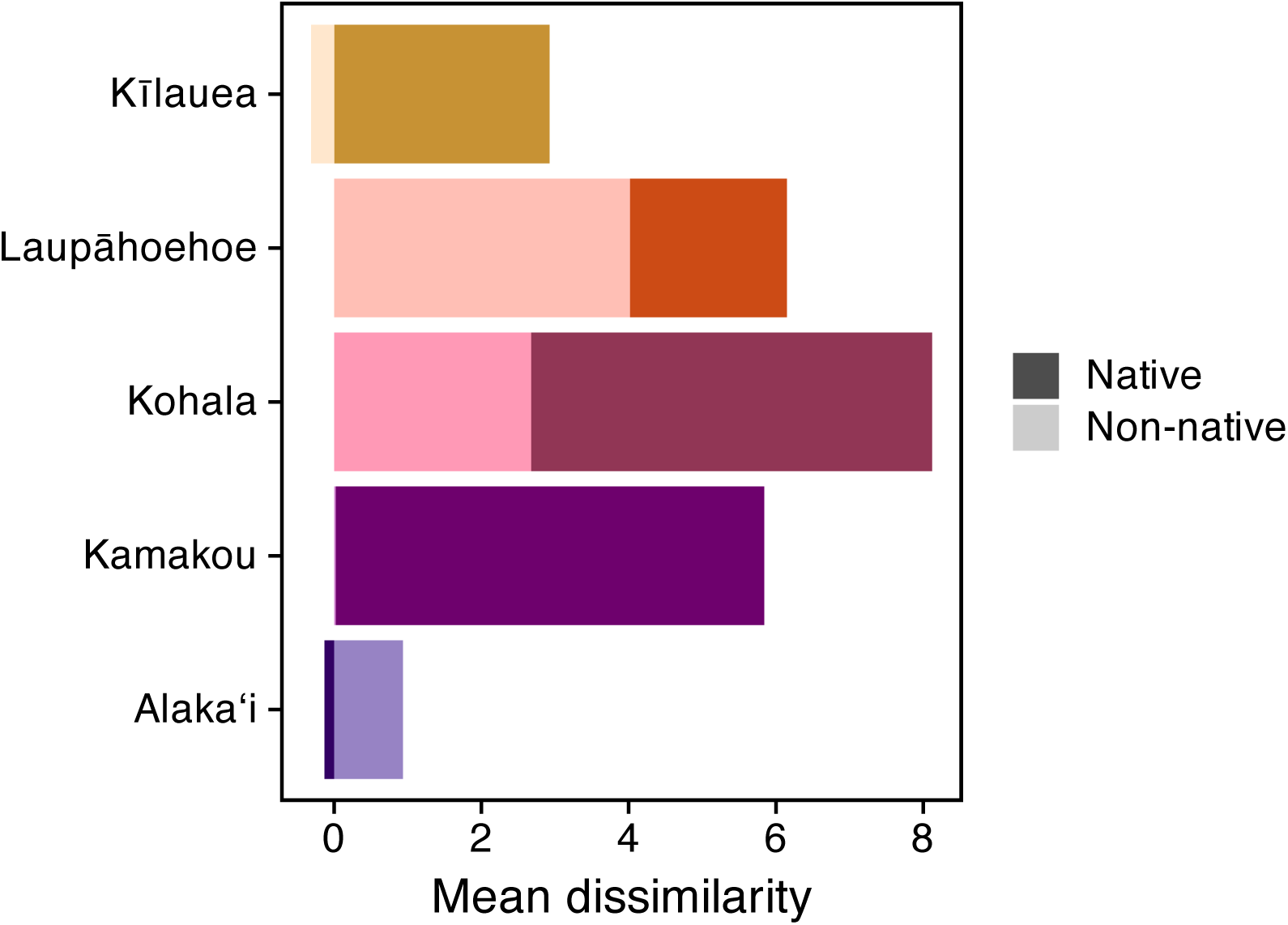
Difference in contribution of native and non-native species to *β*-diversity compared to null. The total height of bars (taking into account negative heights as well) is equal to the overall *β*-diversity z-score for a site. These overall *β*-diversity values are same in pattern, but different in exact value, to those in Figure 5 because the null model used here is more restrictive (see methods). Lighter colors represent the contribution of non-native species and darker colors the contributions of native species to overall *β*-diversity. The contribution of non-native species at Kamakou is small and positive (0.02). The contribution of non-native species at Kīlauea is negative and the contribution of native species at Alaka‘i is negative.

The oldest (Alaka‘i) and youngest (Kīlauea) sites contrast with Laupāhoehoe, Kohala, and Kamakou sites where both native and non-native species make positive contributions to *β*-diversity. In Kohala and Kamakou the positive contribution of native species is substantially larger than non-natives, whereas in Laupāhoehoe the contribution of non-natives is larger.

## Discussion

Real data deviate from both the METE-predicted SAD and IPD in consistent ways across the chronosequence Figure 2. This is strong (sensu McGill 2003) evidence of meaningful departures from the steady state assumptions of METE due to biological processes not accounted for in the deliberately simple state variables of the theory.

Patterns of deviation in the IPD are more variable than those of the SAD. This is not surprising given that the estimation of metabolic rate from allometric relationships introduces more uncertainty than the direct observation of abundances in samples. The choice to use random samples from allometric relationships, rather than expected values, slightly reduces the variability of IPD patterns because expected values introduce artificial discretization of metabolic rate estimates that are inconsistent with METE and indeed the reality that metabolic rates are continuous (see Supplementary Section S6 and the appendix “Sampling biomass estimates from allometric equations”). Nonetheless, rerunning all IPD analyses using expected values from allometric relationships does not substantially change our findings (Supplementary Section S6).

Deviations from the IPD are not consistent in any one direction. Theory both under– and over-predicts the shape of the tails of the IPD in roughly equal proportion. Conversely, deviations in the SAD are consistently in the direction of theory under-predicting unevenness Figure 4. In one way, this result is surprising given that METE predicts a log-series distribution (Harte 2011) which is already one of the most uneven theoretical SAD shapes. However, recent extensive investigation of empirical SADs has found that deviations from statistical baselines akin to METE are often in the direction of more unevenness (Diaz *et al*. 2021). As pointed out in Diaz *et al*. (2021), deviations toward more unevenness could be accounted for by processes that increase the prevalence of rare species, and/or that increase the abundance of common species. Both niche-based (Chesson 2000; Yenni *et al*. 2012) and stochastic dispersal-based (Gotelli 1991) mechanisms could be valid causes of such processes. Complementary to those possible mechanisms is the possibility that adaptive radiation or elevated neutral speciation could produce both many rare species—young and incipient lineages—and a few highly dominant species—those that have found a successful strategy and/or are more established (Gillespie & Whittaker 2025; Hubbell 2001; Schluter 2000; Stroud & Losos 2016). This possibility becomes relevant in hypothesizing mechanisms for the nature of deviations from METE across the chronosequence.

Across geologic ages, both the proportion of non-native species and *β*-diversity at a site predict deviations from METE, but in subtly different yet important ways. Across four out of five sites (all but the geologically youngest) deviations from METE increase with the proportion of non-native species Figure 5. The geologically youngest site, Kīlauea, breaks this pattern with a high proportion of non-native species but a small deviation from METE. Across all five sites deviations from METE are consistently predicted by within site *β*-diversity Figure 5. Both *β*-diversity and proportion of non-native species point to instability being a driver of deviation from METE.

The fact that geologic age seems to modulate the effect of proportion of non-native species but not within site β-diversity begs the question of how age, biological invasion, and β-diversity relate to each other. *β*-diversity partitions unequally between native and non-native species across the chronosequence Figure 6. Kīlauea, the youngest site, sees non-native species contributing negatively (compared to the null) to overall β-diversity, leaving native species making the only positive contribution to overall β-diversity. The contributions of native and non-native species to the β-diversity at the oldest site Alaka‘i are reversed. But at both sites overall β-diversity is low. Thus low β-diversity is more consistently associated with low deviation from METE. The high proportion of non-native species at Kīlauea is not consequential because those non-native species are very homogeneously distributed.

The opposite *β*-diversity contributions of native and non-native species at opposite ends of the chronosequence hints at a mechanistic possibility to explain low β-diversity and low deviation from METE. The assembly of the young Kīlauea site could be dominated by dispersal, which would be consistent with its age—young sites, by definition, have not had sufficient time for diversification and must be assembled by dispersal from source populations. Furthermore, if dispersal is high, this would be consistent with low *β*-diversity driven by native species. Conversely, the oldest Alaka‘i site could represent the outcome of > 4 million years of eco-evolutionary assembly resulting in assemblages that are stable and in steady-state due to niche partitioning and maximized fitness within niches (Barabás *et al*. 2018; Chesson 2000; Godoy *et al*. 2014). This would again result in low β-diversity, with newly arriving non-native species mostly contributing to that low *β*-diversity. In between these mechanistic end points, our findings are consistent with the hypothesis that dispersal-assembled communities in an unstable steady state undergo a non-steady state transition via evolutionary diversification to a stable steady state.

Interestingly, the middle aged sites where we see greatest deviation from METE also show high proportions of non-native species, and high *β*-diversity derived from large contributions of both native and non-native species all occur together Figure 5. These are also the locations along the chronosequence that correspond to the typical minimum time for diversification identified by phylogenetic studies (Bennett & O’Grady 2013; Gillespie 2004; Hormiga *et al*. 2003). The spatiotemporal confluence of these patterns and hypothesized processes along the chronosequence suggests that eco-evolutionary non-steady state might represent an opportunity for biodiversity to change toward a steady state. Left free of anthropogenic pressure, that change could be achieved slowly by *in situ* diversification. Or, in the presence of anthropogenic pressure, that change could be achieved rapidly through biological invasion. This could explain biological invasion following disturbance (anthropogenic or otherwise) (Burke & Grime 1996; D’Antonio & Vitousek 1992; Davis *et al*. 2000; Hobbs & Huenneke 1992). It also predicts that the Hawaiian Archipelago and other hotspots of diversification might be particularly prone to biological invasion.

These hypotheses are, however, only suggested by our work and would require further data collection and testing to support. In particular, extensive population genetic study would be needed to confirm the purported history of dispersal at young sites, population adaptation and bifurcation leading to diversification at middle aged sites, and population stability at older sites. New molecular techniques may soon make such study a reality (Manel *et al*. 2025; Weitemier *et al*. 2021).

