## Supplement for "A maximum entropy perspective reveals deviations from steady state during active diversification"

### Supplement to “Eco-evolutionary deviations from steady state across the Hawai‘i chronosequence revealed with a maximum entropy perspective”

This supplement walks through the steps of recreating the entire analysis presented in the main text. It also provides further analysis in support of and to contextualize results in the main text.

#### S1 Computing setup

Analyses are supported by a custom R package *HawaiiArthMETE* (Rominger & Thai 2026) which can be installed from github.

```
devtools::install_github("ecoevomatics/HawaiiArthMETE")
```

The quarto (Allaire *et al.* 2025) document used to generate this supplement can be found with the R command `system.file("ms/hawaii-arth-mete-supp.qmd", package = "HawaiiArthMETE")`.

The following R computing environment is used for all analyses:

```
library(meteR)
library(ggplot2)
library(ggh4x)
library(dplyr)
```

```
Attaching package: 'dplyr'
```

```
The following objects are masked from 'package:stats':

    filter, lag

The following objects are masked from 'package:base':

    intersect, setdiff, setequal, union
```

```r
library(tidyr)
library(metafor)
```

```
Loading required package: Matrix


Attaching package: 'Matrix'

The following objects are masked from 'package:tidyr':

    expand, pack, unpack

Loading required package: metadat

Loading required package: numDeriv


Loading the 'metafor' package (version 5.0-1). For an
introduction to the package please type: help(metafor)
```

```r
library(HawaiiArthMETE)
```

```


Attaching package: 'HawaiiArthMETE'

The following object is masked from 'package:meteR':

    arth
```

```r
knitr::opts_chunk$set(
    # default figure dims
    fig.width = 5.5, fig.height = 4,
    # caching is default for all subsequent chunks
    cache = TRUE
)

sessionInfo() |>
    print(locale = FALSE)
```

```
R version 4.5.2 (2025-10-31)
Platform: aarch64-apple-darwin20
Running under: macOS Sequoia 15.0.1

Matrix products: default
BLAS:   /System/Library/Frameworks/Accelerate.framework/Versions/A/Frameworks/vecLib.framewo[illegible]
LAPACK: /Library/Frameworks/R.framework/Versions/4.5-arm64/Resources/lib/libRlapack.dylib;  [illegible]

attached base packages:
[1] stats     graphics  grDevices utils     datasets  methods   base

other attached packages:
 [1] HawaiiArthMETE_0.1.0 metafor_5.0-1        numDeriv_2016.8-1.1
 [4] metadat_1.6-0        Matrix_1.7-4         tidyr_1.3.1
 [7] dplyr_1.2.1          ggh4x_0.3.1          ggplot2_4.0.1
[10] meteR_1.2

loaded via a namespace (and not attached):
 [1] gtable_0.3.6       jsonlite_2.0.0     compiler_4.5.2     tidyselect_1.2.1
 [5] scales_1.4.0       yaml_2.3.10        fastmap_1.2.0      lattice_0.22-7
 [9] R6_2.6.1           generics_0.1.4     knitr_1.50         tibble_3.3.0
[13] pillar_1.11.0      RColorBrewer_1.1-3 rlang_1.2.0        mathjaxr_2.0-0
[17] xfun_0.52          S7_0.2.1           cli_3.6.5          withr_3.0.2
[21] magrittr_2.0.4     digest_0.6.39      grid_4.5.2         rstudioapi_0.17.1
[25] nlme_3.1-168       lifecycle_1.0.5    vctrs_0.7.3        evaluate_1.0.4
[29] glue_1.8.0         farver_2.1.2       rmarkdown_2.29     purrr_1.1.0
[33] tools_4.5.2        pkgconfig_2.0.3    htmltools_0.5.8.1
```

The data used in all analyses is available with the *HawaiiArthMETE* package

```r
# load data
data(arth)
```

#### S2 METE calculations

Comparison of the observed data to predictions from the Maximum Entropy Theory of Ecology (METE) is performed with package *meteR* (Rominger & Merow 2017). The following code records information about each sample (name, flow age, proportion non-native presence) as well as several metrics of deviation between theory and observation:

- a $z^2$ value measuring a normalized difference in likelihoods between observed data and theoretical expectation
- differences in proportion of rarity (number of singletons or normalized metabolic rate $\leq 2$) observed and expected
- differences in proportion of dominance (sum of top three most abundant species or sum of top three highest normalized metabolic rates) observed and expected

Note that rarity and dominance metrics differ between SAD and IPD; see Methods of the main text for definitions. All metrics are calculated for both the species abundance distribution (SAD) and individual metabolic rate (or power) distribution (IPD).

```r
arth_mete <- split(arth, arth[, c("site", "tree")], drop = TRUE) |>
    lapply(function(x) {
        # the esf for this grouping
        esf <- meteESF(spp = x$species_code,
                       abund = x$abundance,
                       power = x$metabolic_rate)

        # sad and ipd
        s <- sad(esf)
        p <- ipd(esf)

        data.frame(
            # sample info
            site = x$site[1],
            site_name = x$site_name[1],
            tree = x$tree[1],
            flow_age_my = x$flow_age_my[1],

            # z2 values
            z2_sad = logLikZ(s)$z,
```

```
            z2_ipd = logLikZ(p)$z,

            # differences in extreme values
            n1_diff = log(mean(meteDist2Rank(s) == 1) / mean(s$data == 1)),
            nmax_diff = log(sum(meteDist2Rank(s)[1:10]) / sum(s$data[1:10])),
            p1_diff = log(mean(meteDist2Rank(p) <= 3) / mean(p$data <= 3)),
            pmax_diff = log(sum(meteDist2Rank(p)[1:10]) / sum(p$data[1:10]))
        )
    })

arth_mete <- do.call(rbind, arth_mete)
```

#### S3 Deviations from METE across the chronosequence

Now we analyze how deviations from METE behave across the chronosequence. The following code produces the analysis of a hump-shaped versus flat relationship between METE deviation and flow age.

```
# quadratic model of sad z^2 across age
sad_rma_chrono <- site_means_lm(log(z2_sad) ~
                                    log(flow_age_my) + I(log(flow_age_my)^2) +
                                    (1 | site),
                                data = arth_mete)

# quadratic model of ipd z^2 across age
ipd_rma_chrono <- site_means_lm(log(z2_ipd) ~
                                    log(flow_age_my) + I(log(flow_age_my)^2) +
                                    (1 | site),
                                data = arth_mete)
```

Likelihood ratio tests (LRT) can evaluate the significance of the quadratic model

```
# likelihood ratio tests for sad and ipd across chrono
sad_lrt_chrono <- anova(sad_rma_chrono)
sad_lrt_chrono
```

```
Test of Moderators (coefficients 2:3):
QM(df = 2) = 28.7300, p-val < .0001
```

```r
ipd_lrt_chrono <- anova(ipd_rma_chrono)
ipd_lrt_chrono
```

```
Test of Moderators (coefficients 2:3):
QM(df = 2) = 5.5130, p-val = 0.0635
```

The quadratic model for SAD $z^2$-values is supported by the LRT with a $P$-value $< 0.0001$, but the quadratic model for IPD $z^2$-values just misses the significance threshold of $P \leq 0.05$. However, the Wald-based 95% CI for the quadratic term in the IPD $z^2$-values model does not overlap 0, indicating that by the less stringent Wald-approximation, this quadratic term is significant and the data do tend toward a quadratic shape, just with greater variance compared to the quadratic model for SAD $z^2$-values.

```r
confint(ipd_rma_chrono, fixed = TRUE, random = FALSE)
```

```
                     estimate   ci.lb   ci.ub
intrcpt                0.1040 -0.6281  0.8361
log(flow_age_my)      -0.5985 -1.1176 -0.0793
I(log(flow_age_my)^2) -0.0840 -0.1543 -0.0137
```

The following code produces the visualization reported in the main text of deviation from METE across flow ages.

```r
# prepare data frame of predicted values for plotting
x <- range(arth$flow_age_my) |>
    log() |>
    (\(x) seq(x[1], x[2], length.out = 50))() |>
    exp() |>
    data.frame(flow_age_my = _)

sad_hat <- site_means_predict(sad_rma_chrono, newdata = x)
ipd_hat <- site_means_predict(ipd_rma_chrono, newdata = x)

z2_chrono_pred <- bind_rows(z2_sad = data.frame(x, sad_hat),
                            z2_ipd = data.frame(x, ipd_hat),
                            .id = "metric") |>
    select(!se) |>
    rename(z2 = pred,
```

```r
           z2_lo = ci.lb,
           z2_hi = ci.ub) |>
    mutate(across(starts_with("z2"), exp))


# plotting
fig_z2_chrono <- pivot_longer(arth_mete, c(z2_sad, z2_ipd),
             names_to = "metric", values_to = "z2") |>
    ggplot(aes(flow_age_my, z2, color = site_name)) +
    geom_jitter(width = 0.1) +
    # confidence envelope
    geom_ribbon(data = z2_chrono_pred,
                mapping = aes(x = flow_age_my,
                              ymin = z2_lo,
                              ymax = z2_hi),
                color = "transparent", alpha = 0.4) +
    # central tendency
    geom_line(data = z2_chrono_pred,
              mapping = aes(flow_age_my, z2),
              color = "black") +
    # facet by IPD or SAD and trick strip label into axis label
    facet_grid(
        rows = vars(metric),
        labeller = as_labeller(
            c(z2_ipd = "\"IPD\"~z^2*\"-value\"",
              z2_sad = "\"SAD\"~z^2*\"-value\""),
            label_parsed
        ),
        switch = "y"
    ) +
    xlab("Flow age (my)") +
    this_ms_look(log_x = TRUE, log_y = TRUE) +
    theme(axis.title.y = element_blank(),
          strip.background = element_blank(),
          strip.placement = "outside")

fig_z2_chrono
```

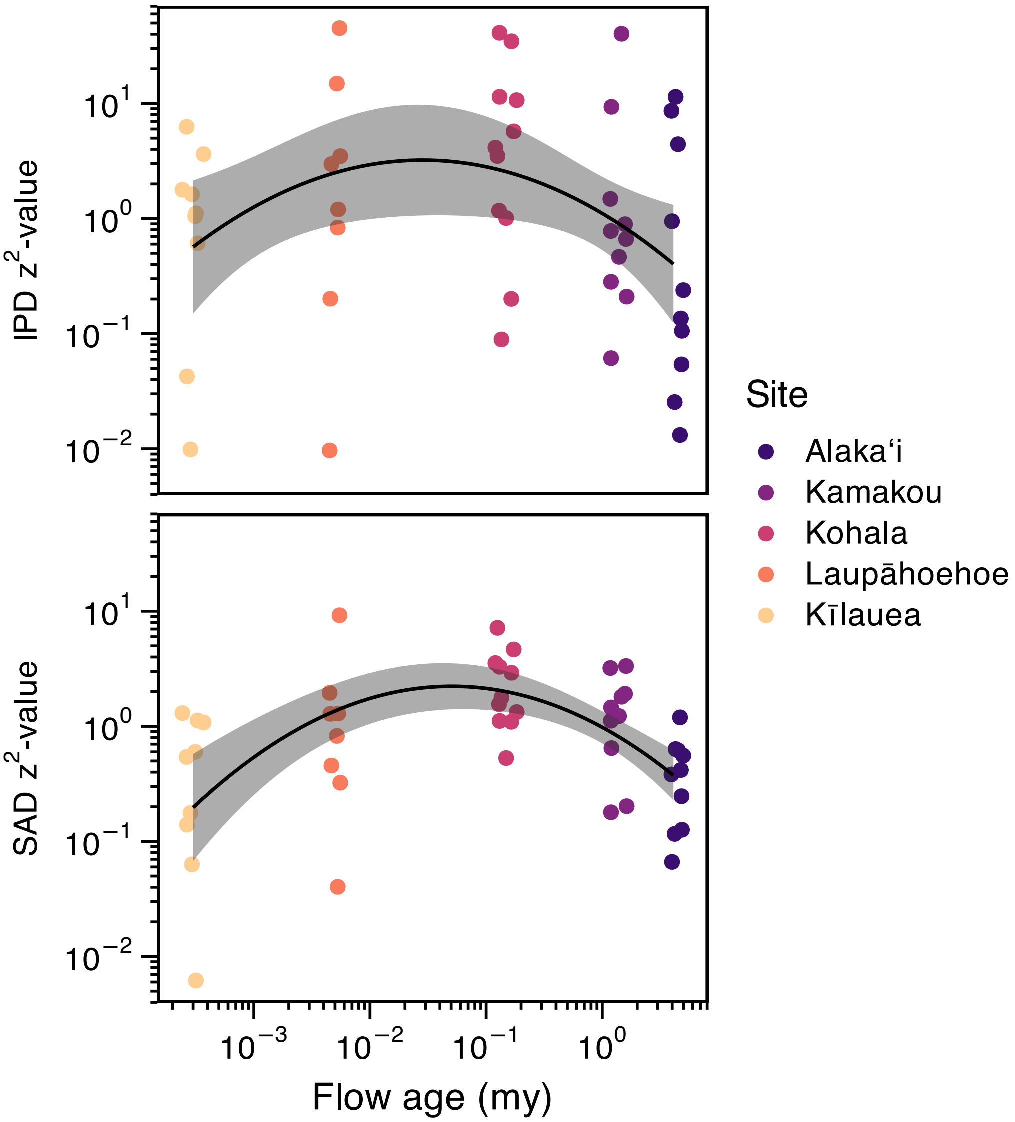


Supplementary Figure S1: Deviations from METE as measured by $z^2$ across the chronosequence.

#### S4 How do observed patterns deviate from METE?

Generally $z^2$-values of the SAD and IPD are correlated Supplementary Figure S2.

```
expl <- data.frame(site = c("VO", "KH"),
                   site_name = c("Kīlauea", "Kohala"),
```

```
                   tree = c(7, 1),
                   deviation = c("lo", "hi"))


ggplot(arth_mete, aes(z2_sad, z2_ipd, color = site_name)) +
    geom_point() +
    geom_point(data = left_join(expl, arth_mete),
               mapping = aes(z2_sad, z2_ipd),
               size = 6,
               show.legend = FALSE) +
    geom_point(data = left_join(expl, arth_mete),
               mapping = aes(z2_sad, z2_ipd),
               size = 6, shape = 1, color = "black",
               stroke = 1.5,
               show.legend = FALSE) +
    xlab(expression("SAD"~z^2*"-values")) +
    ylab(expression("IPD"~z^2*"-values")) +
    this_ms_look(log_x = TRUE, log_y = TRUE)
```

```
Joining with `by = join_by(site, site_name, tree)`
Joining with `by = join_by(site, site_name, tree)`
```

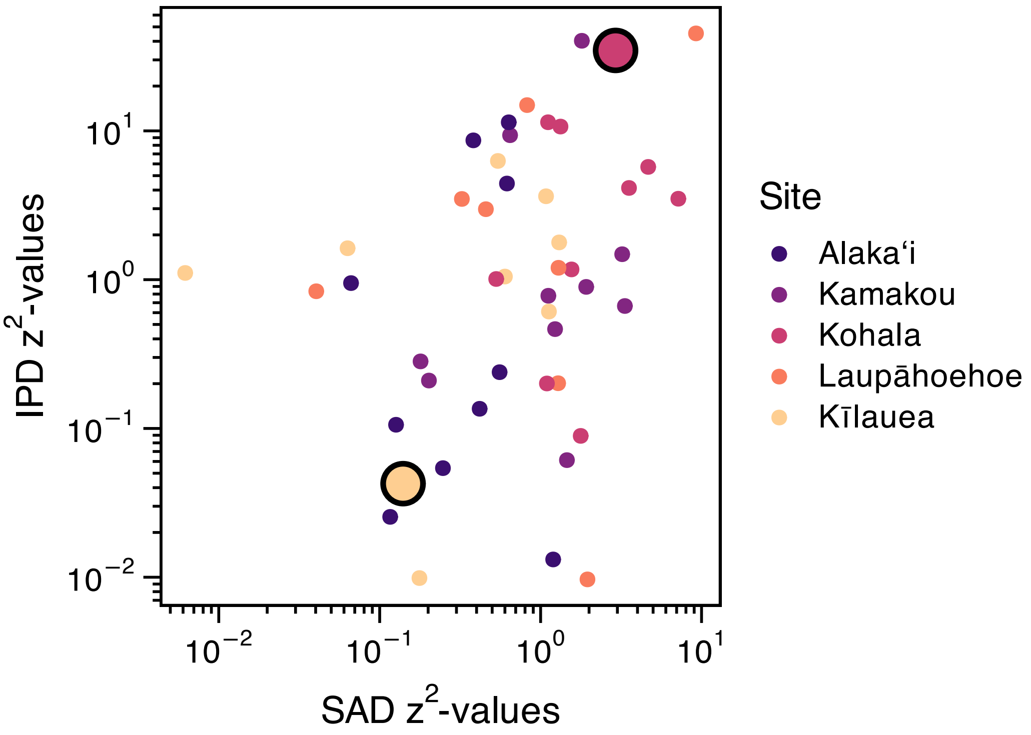


Supplementary Figure S2: Relationship between deviations of SAD and IPD. Slighlty enlarged, circled points are those used as examples of low and high deviation from METE

Next we determine how the observed data deviate from the METE predictions. To begin we look at two exemplar SADs and two exemplar IPDs representing low and high deviation from METE (Supplementary Figure S3). The high and low $z^2$-values of these samples are highlighted in Supplementary Figure S2. Specific samples were chosen such that they have comparable state variables.

```
# function to summarize theoretical and observed distributions
# given one sample `x`

mk_summ <- function(x) {
    esf <- meteESF(x$species_code, x$abundance, x$metabolic_rate)

    list(sad = sad(esf),
         ipd = ipd(esf)) |>
        lapply(function(x) {
            thr <- samp_mete(x, n = 1000) |>
                summarize_samp_mete()
```

```r
            obs <- data.frame(rank = seq_along(x$data),
                              abund_obs = x$data)

            full_join(thr, obs)
        }) |>
        bind_rows(.id = "dist")
}
```

```r
expl_samps <- left_join(expl, arth) |>
    (\(x) split(x, x$deviation, drop = TRUE))() |>
    lapply(mk_summ) |>
    bind_rows(.id = "deviation")
```

```
Joining with `by = join_by(site, site_name, tree)`
Joining with `by = join_by(rank)`
Joining with `by = join_by(rank)`
Joining with `by = join_by(rank)`
Joining with `by = join_by(rank)`
```

```r
fig_how_diff <- ggplot(expl_samps, aes(rank, m)) +
    geom_point(aes(rank, abund_obs)) +
    geom_ribbon(aes(ymin = lo, ymax = hi),
                fill = "#489BC5", alpha = 0.75) +
    geom_line(color = "#489BC5") +
    facet_grid2(
        rows = vars(dist), cols = vars(deviation),
        scales = "free", independent = "x",
        labeller = as_labeller(
            c(hi = "High deviation", lo = "Low deviation",
              ipd = "Metabolic rate", sad = "Abundance")
        ),
        switch = "y"
    ) +
    xlab("Rank") +
    ylab("") +
    this_ms_look(log_y = TRUE) +
    theme(strip.background.y = element_blank(),
          strip.placement.y = "outside",
```

```r
          strip.text.y = element_text(size = 14))

fig_how_diff
```

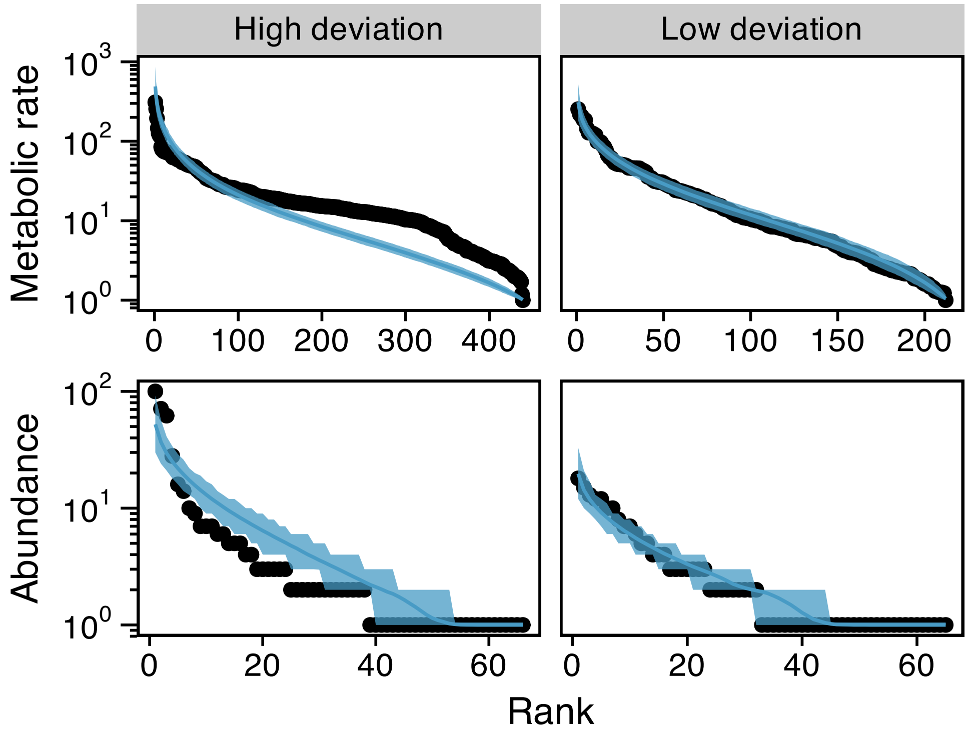


Supplementary Figure S3: Two exemplar IPDs and SADs showing high and low deviation from METE. The low deviation distributions come from the same sample at the Alakaʻi site while the high deviation distributions come from the same sample at the Kamakou site. Samples were choosen to represent ranges of deviation while still being similar in state variables

Looking across all samples (Supplementary Figure S4) we see that deviations between observed and theoretical SADs is consistently in the direction of under-predicted both rarity and dominance (i.e. observed SADs have more rarity and greater dominance than theoretical predictions). Indeed, only one sample shows a (small) over-prediction in dominance, and only two samples show a (small) over-prediction in rarity. This means that observed SADs are more uneven than the METE prediction.

```r
fig_how_z2 <- bind_rows(sad = select(arth_mete, site_name,
                       z2 = z2_sad, rar_diff = n1_diff, dom_diff = nmax_diff),
```

```
          ipd = select(arth_mete, site_name,
                       z2 = z2_ipd, rar_diff = p1_diff, dom_diff = pmax_diff),
          .id = "dist") |>
    pivot_longer(ends_with("diff"),
                 names_to = c("comp", ".value"),
                 names_sep = "_") |>
    ggplot(aes(diff, z2, color = site_name)) +
    geom_point() +
    facet_grid2(rows = vars(dist), cols = vars(comp),
                scales = "free", independent = "x",
                labeller = as_labeller(
                    c(ipd = "\"IPD\"~z^2*\"-value\"",
                      sad = "\"SAD\"~z^2*\"-value\"",
                      dom = "\"Diff. in dominance\"",
                      rar = "\"Diff. in rarity\""),
                    label_parsed
                ),
                switch = "both") +
    xlab("") +
    ylab("") +
    this_ms_look(log_y = TRUE) +
    geom_vline(xintercept = 0, color = "gray") +
    theme(strip.background = element_blank(),
          strip.placement = "outside",
          strip.text.y = element_text(size = 14))


fig_how_z2
```

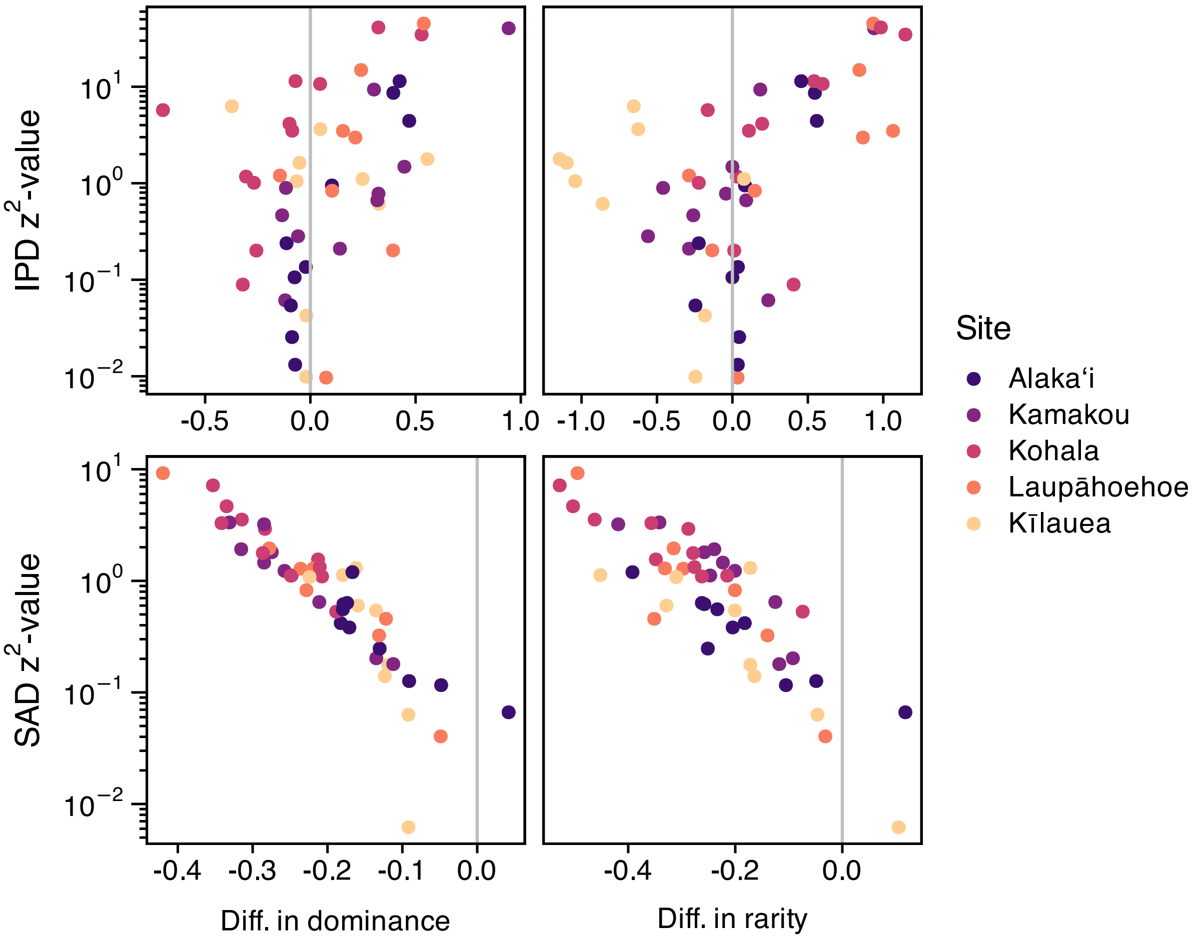


Supplementary Figure S4: Relationship between $z^2$ measure of deviation from METE and different metrics describing how observed data and METE predictions diverge.

In the case of the IPD dominance refers to metabolic rates of individuals with the highest metabolic rates, and rarity refers to the number of individuals with very low metabolic rates. Actual and theoretical IPDs do not show this consistent discrepancy between observed and predicted dominance and rarity. To evaluate this we model the probability of whether the log ratio is above or below zero with an intercept only binomial GLM.

```
# "successes" and "failures" for GLM
ipd_diff <- summarize(arth_mete,
                      n_over_dom = sum(pmax_diff < 0),
                      n_undr_dom = sum(pmax_diff > 0),
                      n_over_rar = sum(p1_diff < 0),
                      n_undr_rar = sum(p1_diff > 0),
```

```r
                      .by = site)

# dominance
pmax_mod <- glm(cbind(n_over_dom, n_undr_dom) ~ 1,
                family = binomial, data = ipd_diff)

summary(pmax_mod)
```

```

Call:
glm(formula = cbind(n_over_dom, n_undr_dom) ~ 1, family = binomial,
    data = ipd_diff)

Coefficients:
             Estimate Std. Error z value Pr(>|z|)
(Intercept) 5.320e-15  2.887e-01       0        1

(Dispersion parameter for binomial family taken to be 1)

    Null deviance: 8.3371  on 4  degrees of freedom
Residual deviance: 8.3371  on 4  degrees of freedom
AIC: 23.11

Number of Fisher Scoring iterations: 3
```

```r
# rarity
p1_mod <- glm(cbind(n_over_rar, n_undr_rar) ~ 1,
              family = binomial, data = ipd_diff)

summary(p1_mod)
```

```

Call:
glm(formula = cbind(n_over_rar, n_undr_rar) ~ 1, family = binomial,
    data = ipd_diff)

Coefficients:
            Estimate Std. Error z value Pr(>|z|)
(Intercept)  -0.3514     0.2994  -1.173    0.241

(Dispersion parameter for binomial family taken to be 1)
```

```
    Null deviance: 14.764  on 4  degrees of freedom
Residual deviance: 14.764  on 4  degrees of freedom
AIC: 28.458

Number of Fisher Scoring iterations: 4
```

The fact that in both models the intercept is not significantly different from the null ($p = 0.5$) means that there is not evidence of systematic difference between theoretical prediction and observed IPD.

### S5 Comparing deviations from METE with potential explanatory variables

#### S5.1 Deviation vs. proportion of non-native species

To see if proportion of non-native species has any relationship with deviation from METE we use meta regression as with the chronosequence analysis. The below code produces this meta regression and tests the affect of proportion of non-native species on deviations of the SAD and IPD from METE predictions.

```
# calculate proportion of non-native spp per site
mete_by_non <- summarize(
    arth,
    prop_nspp_non_nat = n_distinct(
        species_code[!(origin %in% c("endemic", "indigneous"))]) /
        n_distinct(species_code),
    .by = site_name) |>
    # join to mete summary statistics
    full_join(arth_mete)
```

```
Joining with `by = join_by(site_name)`
```

```
# sad regression
sad_rma_non <- site_means_lm(log(z2_sad) ~ prop_nspp_non_nat + (1 | site),
                             # excluding Kīlauea site because it is
                             # quite different, further assessed below
                             filter(mete_by_non, site != "VO"))

# sad likelihood ratio test
```

```r
sad_lrt_non <- anova(sad_rma_non)
sad_lrt_non
```

```
Test of Moderators (coefficient 2):
QM(df = 1) = 22.3287, p-val < .0001
```

```r
# ipd regression
ipd_rma_non <- site_means_lm(log(z2_ipd) ~ prop_nspp_non_nat + (1 | site),
                             # excluding Kīlauea site
                             filter(mete_by_non, site != "VO"))

# ipd likelihood ratio test
ipd_lrt_non <- anova(ipd_rma_non)
ipd_lrt_non
```

```
Test of Moderators (coefficient 2):
QM(df = 1) = 5.2616, p-val = 0.0218
```

The below code is used for visualizing the results. Note that Kīlauea is considered separately because it falls clearly below the expected trend based on other sites. In the case of the SAD, the mean and 95% CI of the $z^2$-value for Kīlauea is below both the expected value and 95% CI of the rest of the data. In the case of the Kīlauea IPD only the mean of $z^2$-values is below the 95% CI of the rest of the data; the two CIs do overlap. This is why in the meta regression analysis Kīlauea was excluded, so we could interrogate the way it deviates from the overall pattern.

```r
# 95% CI for mean of Kīlauea site
vo_mean <- filter(mete_by_non, site == "VO") |>
    pivot_longer(starts_with("z2"),
                 names_to = c(".value", "dist"), names_sep = "_") |>
    mutate(z2 = log(z2)) |>
    summarize(site_name = first(site_name),
              prop_nspp_non_nat = mean(prop_nspp_non_nat),
              z2_mean = mean(z2),
              z2_se = sd(z2) / sqrt(n()),
              z2_lo = z2_mean - 1.96 * z2_se,
              z2_hi = z2_mean + 1.96 * z2_se,
              .by = dist) |>
```

```r
    rename(z2 = z2_mean) |>
    mutate(across(starts_with("z2"), exp))

# prediction for plotting
x <- range(mete_by_non$prop_nspp_non_nat) |>
    (\(x) seq(x[1], x[2], length.out = 50))() |>
    data.frame(prop_nspp_non_nat = _)

sad_hat <- site_means_predict(sad_rma_non, newdata = x)
ipd_hat <- site_means_predict(ipd_rma_non, newdata = x)

z2_non_pred <- bind_rows(sad = data.frame(x, sad_hat),
                         ipd = data.frame(x, ipd_hat),
                         .id = "dist") |>
    select(!se) |>
    rename(z2 = pred,
           z2_lo = ci.lb,
           z2_hi = ci.ub) |>
    mutate(across(starts_with("z2"), exp))

# plotting
fig_dev_non <- pivot_longer(mete_by_non, starts_with("z2"),
                            names_to = c(".value", "dist"),
                            names_sep = "_") |>
    ggplot(aes(prop_nspp_non_nat, z2, color = site_name)) +
    geom_jitter(width = 0.002) +
    # confidence envelope
    geom_ribbon(data = z2_non_pred,
                mapping = aes(x = prop_nspp_non_nat,
                              ymin = z2_lo,
                              ymax = z2_hi),
                color = "transparent", alpha = 0.4) +
    # central tendency
    geom_line(data = z2_non_pred,
              mapping = aes(prop_nspp_non_nat, z2),
              color = "black") +
    geom_rect(data = vo_mean,
              mapping = aes(xmin = prop_nspp_non_nat - 0.002,
                            xmax = prop_nspp_non_nat + 0.002,
                            ymin = z2_lo,
                            ymax = z2_hi),
              color = "transparent", alpha = 0.4) +
```

```r
    geom_segment(data = vo_mean,
                 mapping = aes(x = prop_nspp_non_nat - 0.002,
                               xend = prop_nspp_non_nat + 0.002,
                               y = z2,
                               yend = z2),
                 color = "black") +
    facet_grid(rows = vars(dist),
               labeller = as_labeller(
                   c(ipd = "\"IPD\"~z^2*\"-value\"",
                     sad = "\"SAD\"~z^2*\"-value\""),
                   label_parsed
               ),
               switch = "y") +
    xlab("Proportion of non-native spp.") +
    this_ms_look(log_y = TRUE) +
    theme(axis.title.y = element_blank(),
          strip.background = element_blank(),
          strip.placement = "outside")



fig_dev_non
```

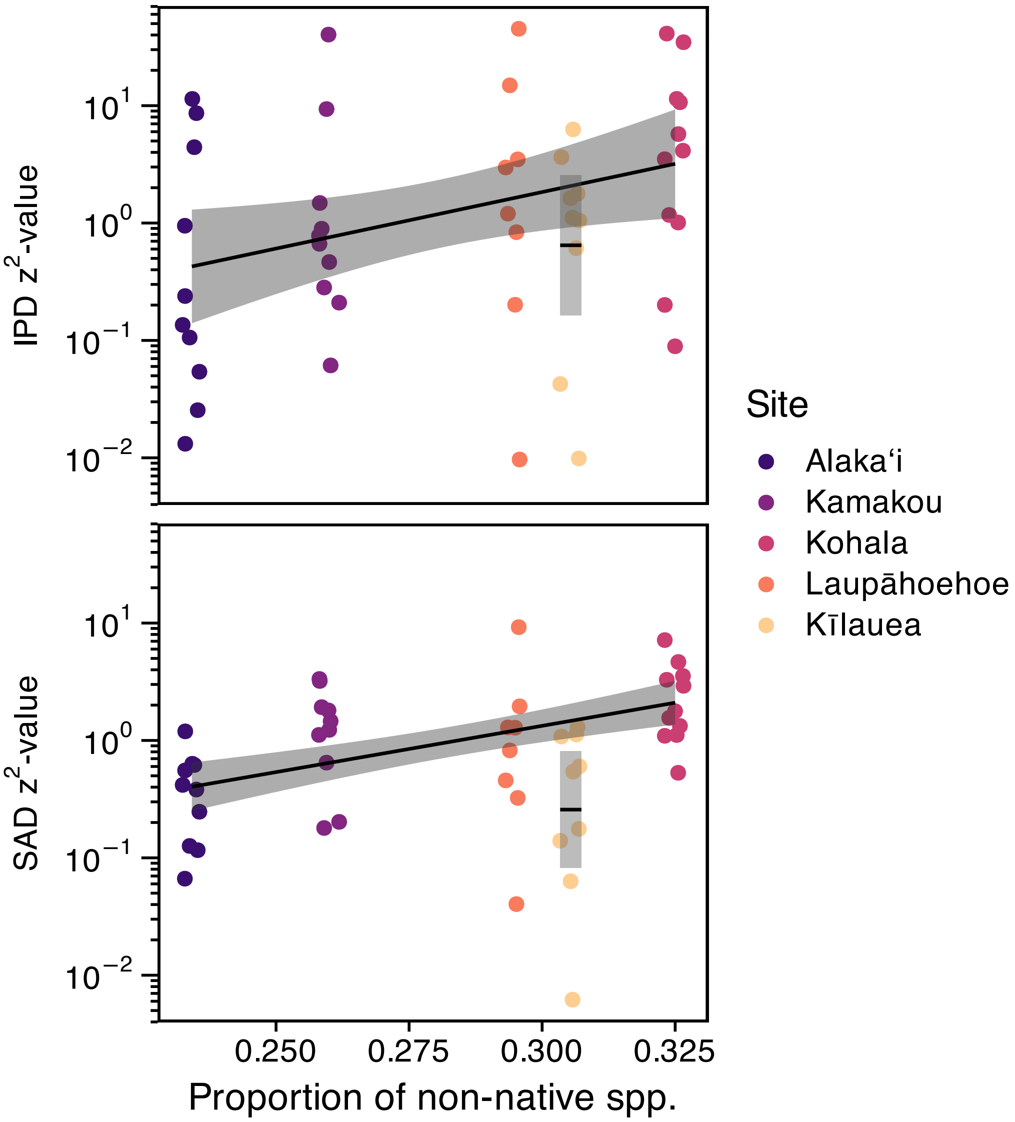


Supplementary Figure S5: Relationship between proportion of non-native species and deviation from METE. Kīlauea was excluded from the overall analysis to be able to compare it separately to the overall trend. Gray regeions represent 95% confidence envelopes (or confidence interval for Kīlauea) and solid lines are expected values.

##### S5.2 Deviation vs. dissimilarity

We also conducted meta regression to see if $z^2$-values change meaningfully across different levels of within-site, between-sample dissimilarity or $\beta$-diversity. The below code runs this meta regression analysis.

```r
# calculate dissimilarity z-scores
diss <- split(arth, arth$site) |>
    sapply(FUN = function(dat) {
        summarize(dat, abund = n(), .by = c(tree, species_code)) |>
            pivot_wider(names_from = species_code,
                        values_from = abund,
                        values_fill = 0) |>
            select(!tree) |>
            bray_diss_z()
    })

# combine with mete summary statistics
mete_by_diss <- mutate(arth_mete,
                       mean_diss = diss[site])



# sad regression
sad_rma_diss <- site_means_lm(log(z2_sad) ~ mean_diss + (1 | site),
                              data = mete_by_diss)

# sad likelihood ratio test
sad_lrt_diss <- anova(sad_rma_diss)
sad_lrt_diss
```

```
Test of Moderators (coefficient 2):
QM(df = 1) = 29.9678, p-val < .0001
```

```r
# ipd regression
ipd_rma_diss <- site_means_lm(log(z2_ipd) ~ mean_diss + (1 | site),
                              data = mete_by_diss)

# ipd likelihood ratio test
ipd_lrt_diss <- anova(ipd_rma_diss)
ipd_lrt_diss
```

```
Test of Moderators (coefficient 2):
QM(df = 1) = 5.4708, p-val = 0.0193
```

The relationship between deviation from METE and dissimilarity is more clearly linear than that of proportion non-native species and deviation from METE. It is therefore worth figuring out if the ways that native and non-native taxa contribute to dissimilarity is different across sites.

The below code visualizes the results of the meta regression or SAD and IPD $z^2$-values against dissimilarity.

```
# prediction for plotting
x <- range(mete_by_diss$mean_diss) |>
    (\(x) seq(x[1], x[2], length.out = 50))() |>
    data.frame(mean_diss = _)

sad_hat <- site_means_predict(sad_rma_diss, newdata = x)
ipd_hat <- site_means_predict(ipd_rma_diss, newdata = x)

z2_diss_pred <- bind_rows(sad = data.frame(x, sad_hat),
                          ipd = data.frame(x, ipd_hat),
                          .id = "dist") |>
    select(!se) |>
    rename(z2 = pred,
           z2_lo = ci.lb,
           z2_hi = ci.ub) |>
    mutate(across(starts_with("z2"), exp))

# plotting
fig_diss_z <- pivot_longer(mete_by_diss, starts_with("z2"),
                           names_to = c(".value", "dist"),
                           names_sep = "_") |>
    ggplot(aes(mean_diss, z2, color = site_name)) +
    geom_jitter(width = 0.1) +
    # confidence envelope
    geom_ribbon(data = z2_diss_pred,
                mapping = aes(x = mean_diss,
                              ymin = z2_lo,
                              ymax = z2_hi),
                color = "transparent", alpha = 0.4) +
    # central tendency
    geom_line(data = z2_diss_pred,
              mapping = aes(mean_diss, z2),
              color = "black") +
    facet_grid(rows = vars(dist),
               labeller = as_labeller(
```

```
                   c(ipd = "\"IPD\"~z^2*\"-value\"",
                     sad = "\"SAD\"~z^2*\"-value\""),
                   label_parsed
               ),
               switch = "y") +
    xlab("Mean dissimilarity") +
    this_ms_look(log_y = TRUE) +
    theme(axis.title.y = element_blank(),
          strip.background = element_blank(),
          strip.placement = "outside")

fig_diss_z
```

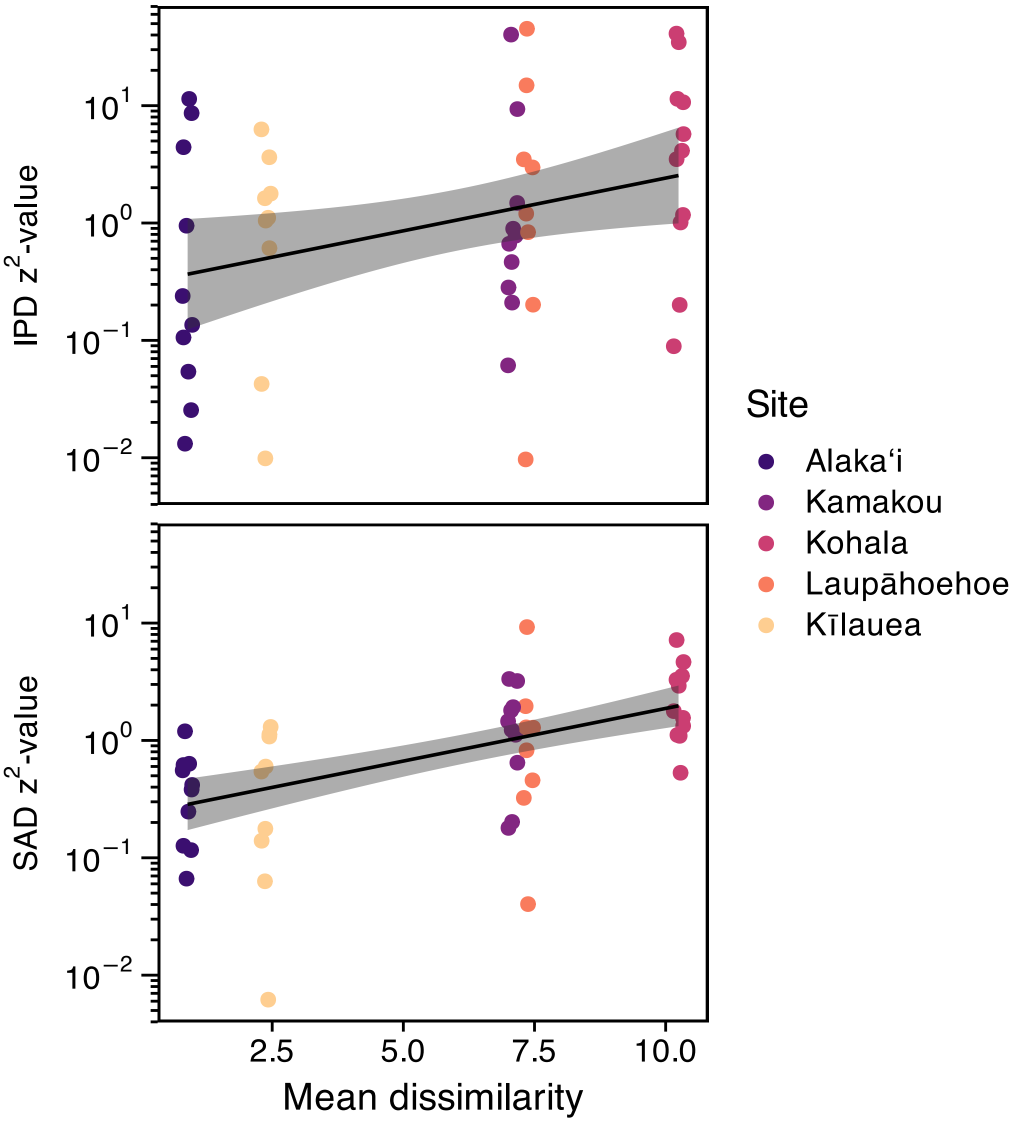


Supplementary Figure S6: Relationship between dissimilarity and deviation from METE. Gray regreions represent 95% confidence envelopes and solid lines are expected values.

##### S5.3 Partitioning Bray Curtis dissimilarity between native and non-native species

To do that we take advantage of the additive nature of Bray Curtis dissimilarity. Bray Curtis dissimilarity is defined as

$$d_{kj} = \frac{\sum_i |x_{ij} - x_{ik}|}{\sum_{ij} x_{ij} + x_{ik}}$$

where $x_{ij}$ and $x_{ik}$ are the abundances of species $i$ at sites $j$ and $k$. This can be partitioned across any grouping of species, say $(a)$ and $(b)$ with abundances $x_{ij}^{(a)}$ and $x_{ij}^{(b)}$

$$d_{kj} = \frac{\sum_i |x_{ij}^{(a)} - x_{ik}^{(a)}|}{\sum_{ij} x_{ij} + x_{ik}} + \frac{\sum_i |x_{ij}^{(b)} - x_{ik}^{(b)}|}{\sum_{ij} x_{ij} + x_{ik}}$$

Now we can look separately at the contribution of group $(a)$ and group $(b)$ to the overall dissimilarity. We use this approach to compare the contributions of native and non-native taxa to overall dissimilarity. We again make use of the `quasiswap_count` permutational null model to standardize these contributions. However, we must permute the two submatrices (one for native species, one for non-natives) separately in order to preserve the distribution of native and non-native individuals across sites. We can then combine these separately permuted submatrices and partition dissimilarity. Comparing observed Bray Curtis partitions to their null counterparts gives us a z-score standardized view of how native and non-native species contributed to overall dissimilarity. The custom function `bray_part_z` from *HawaiiArthMETE* calculates these z scores.

```
site_bray_z <- split(arth, arth$site_name) |>
    lapply(function(dat) {
        comm <- summarize(dat,
                          abund = n(),
                          .by = c(tree, species_code)) |>
            pivot_wider(names_from = species_code,
                        values_from = abund,
                        values_fill = 0) |>
            select(!tree)
        g <- dat$origin[match(names(comm), dat$species_code)] %in%
            c("endemic", "indigenous")

        bp <- bray_part_z(as.matrix(comm), g)
        data.frame(bp_nat = bp[1], bp_non = bp[2])
    }) |>
    bind_rows(.id = "site_name")

# make site names factors ordered by age
site_bray_z$site_name <- factor(site_bray_z$site_name,
                                levels = levels(arth$site_name))
```

Now we can visualize different contributions of native and non-native species to within-site, between-sample dissimilarity.

```
# palette for native and non-native by site
pal <- this_ms_look()[[1]]$palette(5)
pal <- rbind(colorspace::darken(pal, amount = 0.25),
             colorspace::lighten(pal, amount = 0.5)) |>
    as.vector()

fig_bray_part <- site_bray_z |>
    pivot_longer(!site_name) |>
    mutate(g = paste(as.numeric(site_name),
                     name, sep = "_")) |>
    # plotting
    ggplot(aes(value, site_name, fill = g, alpha = name)) +
    geom_bar(stat = "identity") +
    this_ms_look() +
    scale_fill_manual(values = pal, guide = "none") +
    scale_alpha_manual(
    values = c(bp_nat = 1, bp_non = 1), # over-ride alpha
    name = NULL,
    labels = c(bp_nat = "Native", bp_non = "Non-native"),
    guide = guide_legend(override.aes = list(fill = c("grey30", "grey80"),
                                             alpha = 1))) +
    xlab("Mean dissimilarity") +
    theme(axis.title.y = element_blank())
```

```
Scale for fill is already present.
Adding another scale for fill, which will replace the existing scale.
```

```
fig_bray_part
```

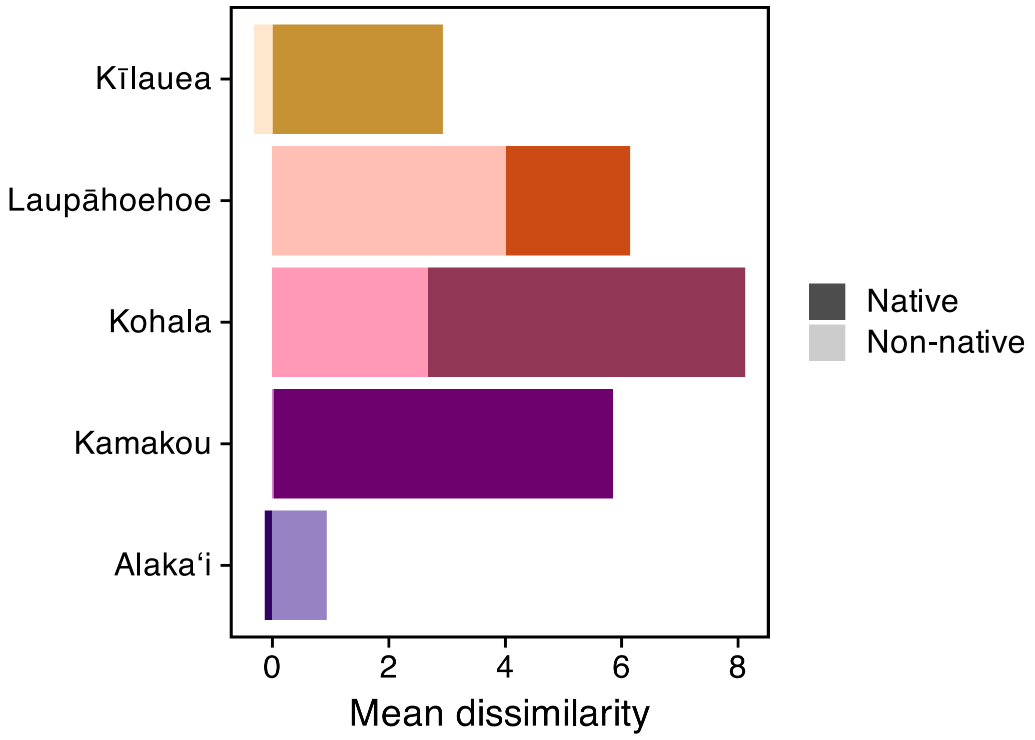


Supplementary Figure S7: Difference in contribution of native and non-native species to dissimilarity compared to null. Negative values indicate greater contribution of non-native species compared to chance while positive values indicate greater contribution of native species compared to chance.

#### S6 Evaluating the consequences of sampling biomass values

Here we re-run all analyses using a single expected value of biomass derived from taxon- and stage-specific allometric scaling relationships rather than sampling with noise from those relationships. To do so we extract all R code from the qmd file generating this supplement, alter the metabolic rate estimates going into the METE calculations, and re-run the code

```
# create r script from qmd
r_temp <- tempfile(fileext = ".R")
quarto::qmd_to_r_script("hawaii-arth-mete-supp.qmd", r_temp)

# remove all chunks subsequent to bray-part (including bray-part)
# because those do not have to do with IPD
r <- readLines(r_temp)
```

```r
r <- r[-(grep("label: bray-part", r)[1]:length(r))]

# remove chunks before `mete-calc`
r <- r[-(1:(grep("label: mete-calc", r) - 1))]

# change metabolic rate to expected value
r <- gsub("x\\$metabolic_rate",
          "x\\$ind_biom\\^0.75",
          r)

# write and source, sourcing into local environment preserves global
# environment
writeLines(r, r_temp)
scr_env <- new.env()
source(r_temp, local = scr_env)
```

Now that all analyses involving the IPD have been re-run with expected values of biomass rather than random draws, we can evaluate differences between the two approaches.

The first thing to notice is that IPD $z^2$-values are correlated under both analysis scenarios Supplementary Figure S8

```r
data.frame(ipd_draw = arth_mete$z2_ipd,
           ipd_expt = scr_env$arth_mete$z2_ipd) |>
    ggplot(aes(ipd_draw, ipd_expt)) +
    geom_point() +
    this_ms_look(log_x = TRUE, log_y = TRUE) +
    geom_abline(intercept = 0, slope = 1) +
    xlab(expression("Random draw IPD"~z^2)) +
    ylab(expression("Expected value IPD"~z^2))
```

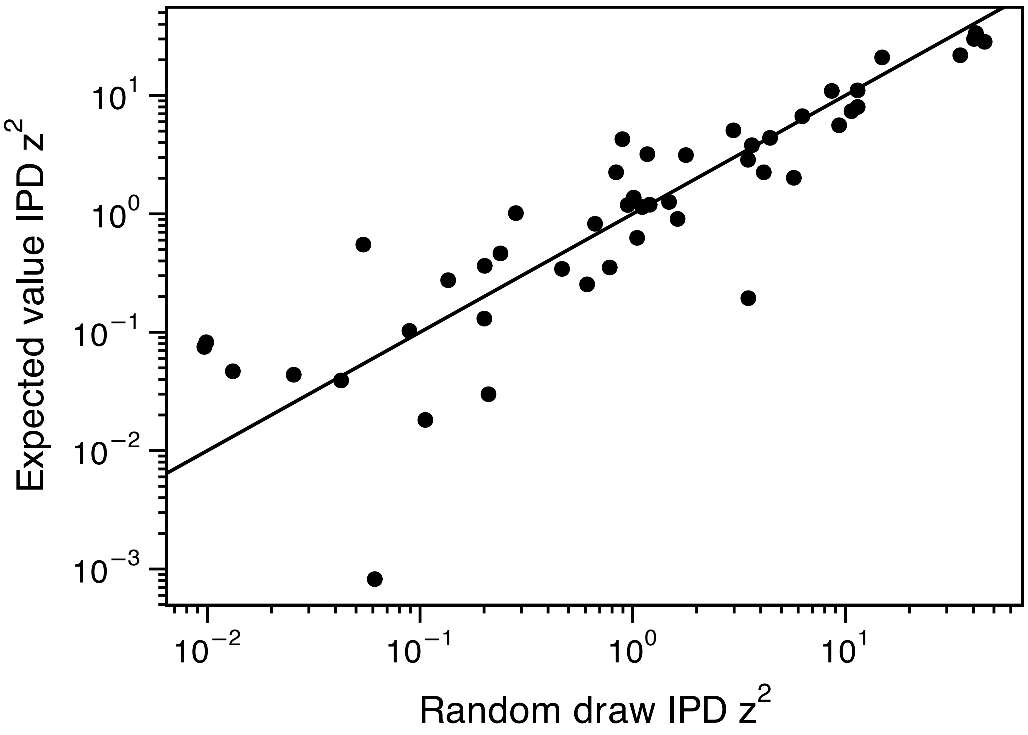


Supplementary Figure S8: Relationship between IPD $z^2$-values when calculated using random draws of expected values of biomass. The solid line is the 1:1 line.

Using expected values for biomass increases the variability of log-transformed IPD $z^2$-values slightly: $SD_{\text{expected values}} = 2.222$ compared to $SD_{\text{random draws}} = 2.236$

This increased variability is not surprising given the artificial descritization that results from using expected values. We can see the impact of this descritization by again comparing the exemplar samples with high and low deviation from Supplementary Figure S2 and Supplementary Figure S3. Now we compare on the IPD $z^2$ values and see that, while the overall shape of the IPD is similar, long runs of the same metabolic rate drive both spurious deviations and conformations to METE predictions Supplementary Figure S9.

```
bind_rows(draws = expl_samps,
          exptd = scr_env$expl_samps,
          .id = "biomass") |>
    filter(dist == "ipd") |>
    ggplot(aes(rank, m)) +
    geom_point(aes(rank, abund_obs)) +
    geom_ribbon(aes(ymin = lo, ymax = hi),
                fill = "#489BC5", alpha = 0.75) +
    geom_line(color = "#489BC5") +
```

```
    facet_grid2(
        rows = vars(deviation), cols = vars(biomass),
        scales = "free", independent = "x",
        labeller = as_labeller(
            c(hi = "High deviation", lo = "Low deviation",
              draws = "Random draws of biomass",
              exptd = "Expected values of biomass")
        )
    ) +
    xlab("Rank") +
    ylab("Metabolic rate") +
    this_ms_look(log_y = TRUE)
```

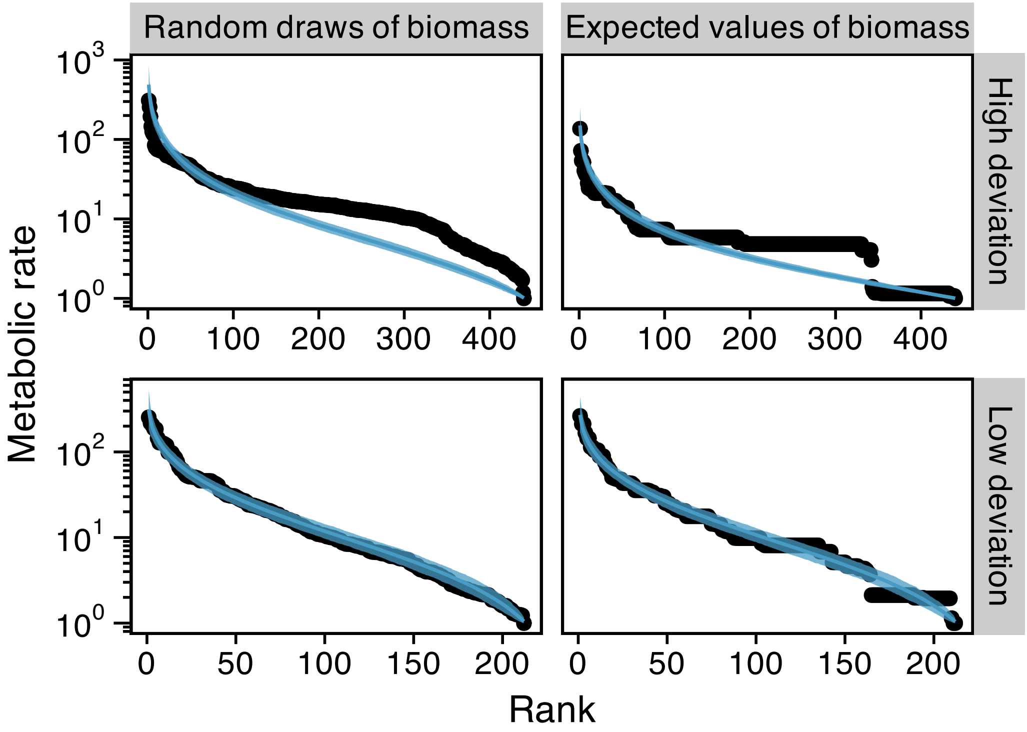


Supplementary Figure S9: Example rank distributions of individual metabolic rate showing both high and low deviations from METE as well as metabolic rate estimates based on random draws from allometric relationships versus the expected values of those relationships. Note that with expected values artificial descritization causes both spurious deviation from METE (as in the low deviation example) and spurios conformations to METE (as in the high deviation example).

The primary effect of increased variability in IPD $z^2$-values and spurious deviations/conformations is to decrease the significance of the relationship of IPD $z^2$-values and explanatory variables. While LRT identified both proportion non-native species and dissimilarity as predictors of IPD $z^2$-values, when using expected values of biomass the $P$-values of these relationships are no longer $\leq 0.05$:

```
# LRT of proportion non-native species
anova(scr_env$ipd_rma_non)
```

```
Test of Moderators (coefficient 2):
QM(df = 1) = 2.8824, p-val = 0.0895
```

```
# LRT of dissimilarity
anova(scr_env$ipd_rma_diss)
```

```
Test of Moderators (coefficient 2):
QM(df = 1) = 2.5978, p-val = 0.1070
```

It should be noted, however, that there is still marginal support for these relationships based on 95% confidence intervals of their coefficients in the two models:

```
# slope of proportion non-native species
confint(scr_env$ipd_rma_non, fixed = TRUE, random = FALSE) |>
    as.data.frame() |>
    (\(x) x[2, , drop = FALSE])()
```

```
                  estimate     ci.lb    ci.ub
prop_nspp_non_nat  16.7525 -2.587085 36.09208
```

```
# slope of dissimilarity
confint(scr_env$ipd_rma_diss, fixed = TRUE, random = FALSE) |>
    as.data.frame() |>
    (\(x) x[2, , drop = FALSE])()
```

```
           estimate       ci.lb     ci.ub
mean_diss 0.1414801 -0.03056349 0.3135237
```

In both cases, the 95% CIs contain a majority of positive values.
